# Efficient and adaptive sensory codes

**DOI:** 10.1101/669200

**Authors:** Wiktor Młynarski, Ann M. Hermundstad

## Abstract

The ability to adapt to changes in stimulus statistics is a hallmark of sensory systems. Here, we develop a theoretical framework that can account for the dynamics of adaptation from an information-processing perspective. We use this framework to optimize and analyze adaptive sensory codes, and we show that codes optimized for stationary environments can suffer from prolonged periods of poor performance when the environment changes. To mitigate the adversarial effects of these environmental changes, sensory systems must navigate tradeoffs between the ability to accurately encode incoming stimuli, and the ability to rapidly detect and adapt to changes in the distribution of these stimuli. We derive families of codes that balance these objectives, and we demonstrate their close match to experimentally-observed neural dynamics during mean and variance adaptation. Our results provide a unifying perspective on adaptation across a range of sensory systems, environments, and sensory tasks.

## INTRODUCTION

The natural sensory environment is constantly in flux. When environmental conditions change, so too do the statistics of sensory stimuli. In response to these changes, sensory systems modify the ways in which incoming stimuli are encoded in neural response patterns. This phenomenon, known as sensory adaptation, has been a subject of research since the beginnings of sensory neuroscience [1].

Adaptation occurs across sensory systems, regardless of the sensory modality or stage along the ascending sensory pathway. Adaptation has been observed in the visual [2], auditory [3–5], somatosensory [6, 7], olfactory [8], and electrosensory [9] systems. It is manifested in individual neurons [10, 11] and neural populations [5, 12, 13], beginning in the sensory periphery [14–16] through the midbrain [17–20] to the cortex [6, 21, 22]. While the most extensively-studied forms of adaptation are induced by changes in low-order stimulus statistics, such as mean [23] or variance [5, 10, 24], sensory systems have also been shown to adapt to more complex statistics, such as the spatial scale [25] and high-order structure [12] of visual stimuli. Despite the prevalence of adaptation across the brain, it remains unclear as to whether this diverse range of phenomena emerges from a common set of neural coding principles.

A dominant hypothesis is that adaptation reflects a dynamic reallocation of finite metabolic resources to support the performance of different sensory tasks [10, 11, 23, 26]. As postulated by the efficient coding hypothesis [27], sensory neurons might be adapted to stimulus statistics in a manner that maximizes the amount of information that they transmit downstream. Different variants of this hypothesis propose that sensory systems might only need to encode specific features of incoming sensory signals, depending on the task at hand [28–30]. Regardless of the task, however, this hypothesis dictates that the neural code must be adapted to the stimulus distribution in order to efficiently encode and transmit task-relevant information. As such, this hypothesis accounts for the *consequences* of sensory adaptation: in order to maintain a given level of task performance, sensory neurons should modify their encoding properties when the distribution of incoming stimuli changes, and efficient coding specifies what these properties should be when the environment is stationary and the sensory code is adapted. It does not, however, specify how the modification of these properties should be accomplished, and whether this modification necessitates different coding strategies than would be required in a stationary setting.

In this work, we extend the efficient coding framework to nonstationary environments and to adaptive sensory representations, and we show that in certain scenarios, adaptive codes must use different strategies than non-adaptive ones. We consider sensory systems that use an internal estimate of the current sensory context, specified by the distribution of incoming stimuli, to guide adaptation [31–33]. When the environment is stationary, this estimate can in principle be used to design a code that maximizes task performance, as posited by classic theories of efficient coding [15, 34, 35]. However, when the environment changes, mismatches between this estimate and the current context can be a substantial source of error. Optimal adaptive codes must therefore devote some of their limited resources to either detecting changes in the underlying distribution of stimuli, or to reducing coding errors induced by an incorrect estimation of the underlying context. We extend the efficient coding framework to formalize this interplay and derive codes that optimally balance these competing objectives. We then use this framework to make and test specific predictions about a well-studied paradigm for characterizing sensory adaptation. Finally, we demonstrate how this framework can serve as a basis for comparing adaptation schemes across a range of different sensory representations, tasks, and environments. Together, these results generalize current theories of neural computation to nonstationary environments, where adaptation is a necessity.

## RESULTS

Consider a sensory neuron that encodes local image features across different visual scenes (Fig 1a). The stochastic response of this neuron can be captured by a linear filter applied to individual image patches, followed by a saturating nonlinearity that transforms noisy filter outputs into Poisson-distributed firing rates [36, 37]. In order to accurately decode the filter outputs, it is well known that the saturating nonlinearity must be adapted to the distribution of these outputs [11, 15, 30, 38, 39].

**Figure 1:**
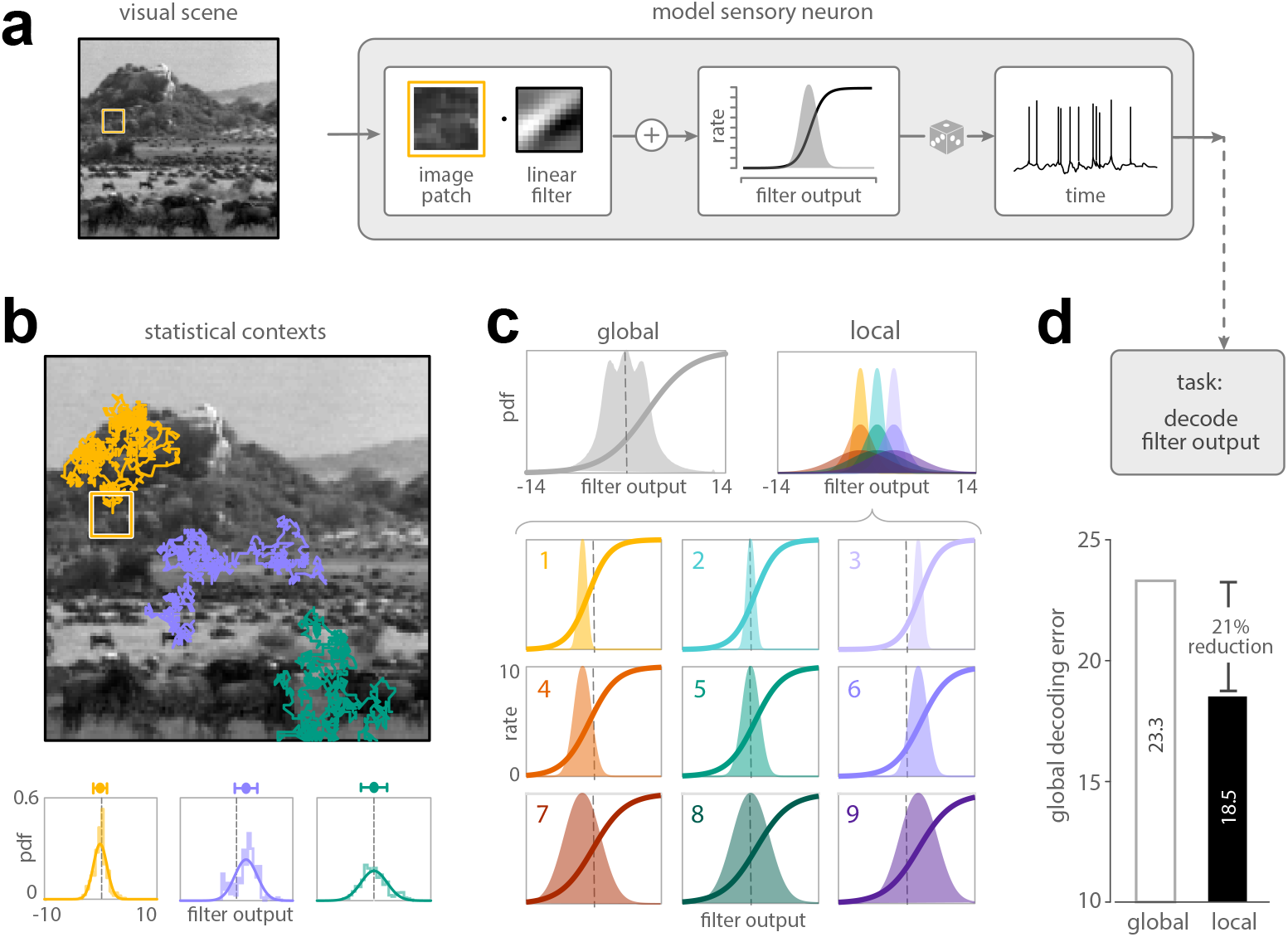
Adaptation can improve performance on sensory tasks. **a)** A linear-nonlinear Poisson neuron encodes images patches from different regions of a visual scene. Local image patches are first filtered with a linear receptive field. The output of this filter is then distorted by noise, passed through a saturating nonlinearity, and used to generate a spiking response via a Poisson process. The saturating nonlinearity can be optimized for the task of decoding the filter output from the spiking response of the neuron. Average decoding error is computed as the mean squared error between the maximum-likelihood estimate and the true value of the filter output. **b)** Visual exploration of different regions of a visual scene (colored lines; main panel) results in different distributions of the filter output (lower panels). **c)** We approximate the variability in filter outputs by a set of nine Gaussian distributions that span a range of means and variances observed across an ensemble of natural scenes. A global nonlinearity (upper left panel) minimizes average decoding error given the distribution of filter outputs marginalized over all nine contexts. Local nonlinearities (lower panels) minimize average decoding error given the distribution of filter outputs within each individual context. **d)** When averaged over filter outputs drawn evenly from each of the nine contexts, the decoding error is higher if the neuron uses a fixed global nonlinearity for all contexts than if it locally adapts the nonlinearity to match each individual context.

During visual exploration of a natural scene, the distribution of these filter outputs will vary (Fig 1b; [40, 41]). This variability can be summarized by a set of distributions that represent different sensory contexts and that span a range of local luminance (mean) and contrast (variance) values observed across an ensemble of natural scenes (Fig. 1c; lower panels).

To minimize decoding error, the nonlinearity might be globally adapted to the marginal distribution of filter outputs averaged across all contexts (Fig 1c; upper left). A more advantageous strategy requires the neuron to locally adapt its nonlinearity to match each individual context (Fig 1c; upper right), such that it devotes its limited dynamic range to stimuli encountered in that context (Fig 1c; lower panels) [10, 11, 23, 41]. This local adaptive strategy reduces decoding error by approximately 21% beyond what is achieved when the same neuron relies on a global nonlinearity adapted to all contexts (Fig 1d).

### A normative framework for adaptive sensory codes

This simple model neuron highlights the well-known advantages of adapting to context-dependent changes in input statistics [11, 41]. However, this form of adaptation assumes that the neuron has perfect knowledge of context that it can use to optimally adapt its nonlinearity. In reality, sensory systems must infer context from incoming sensory stimuli, and can at best use an *estimate* of context to adapt. Because of errors inherent to any estimation process, there will be times when the system is using a suboptimal strategy that is not matched to the current input statistics (Fig 2a). In what follows, we demonstrate that these periods of mismatch can be a considerable source of error in neural coding. We then show how sensory systems can reduce these mismatch-induced errors, and how this in turn can impact performance during times when the system is correctly matched to the input statistics.

**Figure 2:**
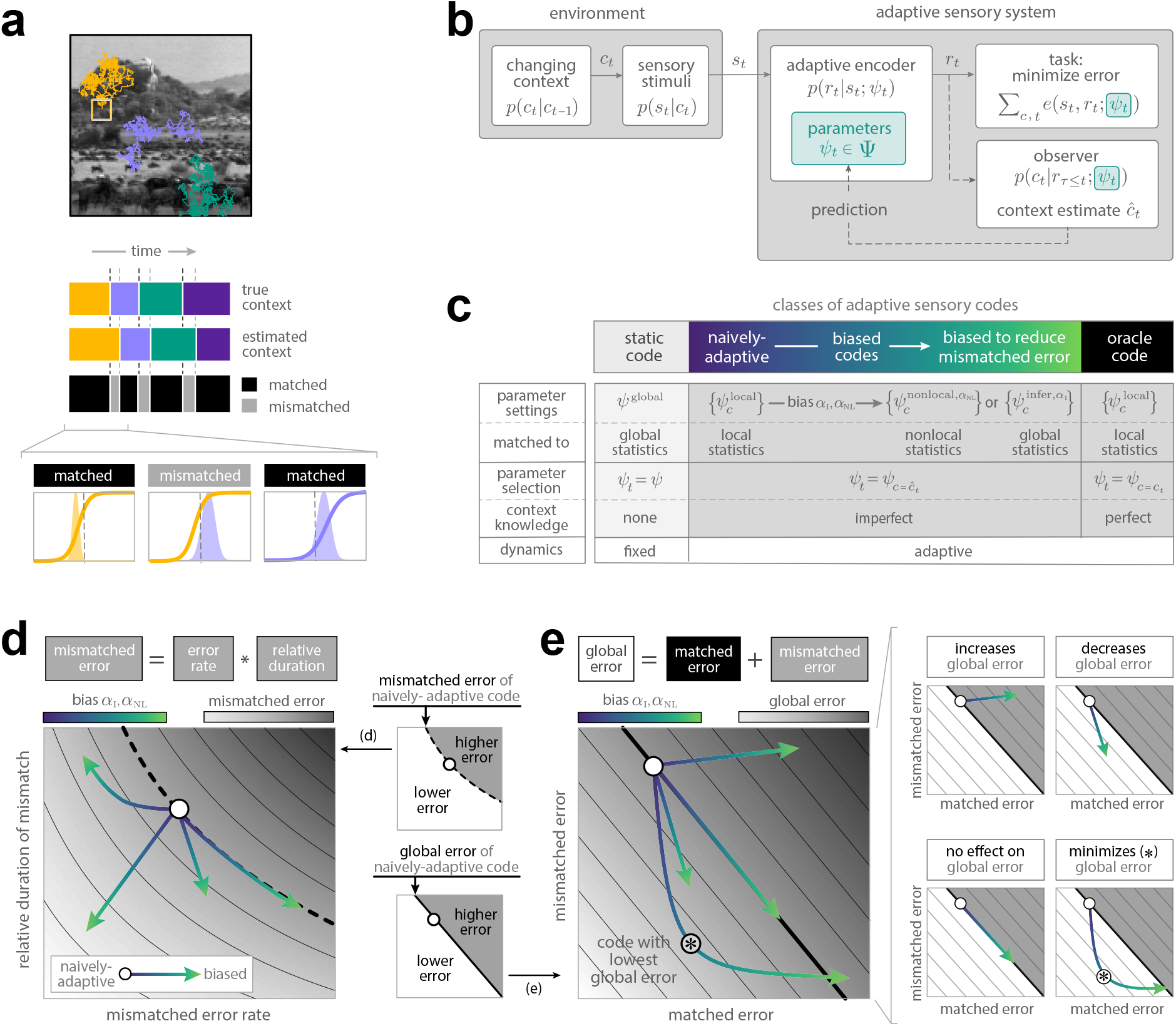
Adaptive sensory codes exhibit different patterns of error. **a)** A sensory system that must estimate a changing context will experience periods of time when this estimate is matched (black) or mismatched (gray) to the current context, illustrated for the example shown in Fig 1. We design adaptive codes that can reduce these periods of mismatch. **b)** We consider a general scenario in which a changing context is signaled by incoming sensory stimuli that are transformed into neural responses by an adaptive encoder with parameters *ψ* (details in text). The output of this encoder is used to perform a sensory task. In order to improve task performance, the encoder can use an internal estimate of context to adapt its encoding parameters. **c)** We organize sensory codes according to whether their encoding parameters are fixed in time or adaptive, and if adaptive, whether they have perfect or imperfect knowledge of context. At one extreme, the “static” code is matched to the distribution of stimuli across contexts; at the other extreme, the “oracle” code is matched to individual contexts. In between, we construct families of biased codes that have imperfect context knowledge and must balance matched and mismatched errors. **d)** Mismatched error (heatmap) is the product between the mismatched error rate (horizontal axis) and the duration of mismatch (vertical axis). We begin with a naively-adaptive code (open circle), and we bias this code toward decreasing either the error rate or the duration of mismatches. The resulting families of biased codes (colored arrows) can occupy different regions of this space. Inset, upper right: the thick dashed line partitions the mismatched error surface into regions that achieve higher (gray) or lower (white) mismatched error than the naively-adaptive code. **e)** The global error (heatmap) is the sum of matched error (horizontal axis) and mismatched error (vertical axis). Inset, lower left: the thick solid line partitions the global error surface into regions that achieve higher (gray) or lower (white) global error than the naively-adaptive code. Biased codes are guaranteed to have higher matched error than the naively-adaptive code, but can have differing impact on global error (four small panels). We seek the code that achieves the lowest global error, marked by the star.

To understand the interplay between these coupled sources of error, we consider an adaptive system that must perform a sensory task in a changing environment (Fig 2b). At a given time *t*, the system receives incoming stimuli *s*_*t*_. The distribution of these stimuli, *s*_*t*_ ∼ *p*(*s*_*t*_|*c*_*t*_), is controlled by a latent context *c*_*t*_ that can change over time, with dynamics governed by *p*(*c*_*t*_|*c*_*t*−1_). The system transforms incoming stimuli into neural responses *r*_*t*_ ∼ *p*(*r*_*t*_|*s*_*t*_; *ψ*_*t*_) using an adaptive encoder that has limited coding precision, and that is controlled by encoding parameters *ψ*_*t*_ selected from a set Ψ. These neural responses are then used to perform a sensory task whose instantaneous error *e*_*t*_(*ψ*_*t*_) ≡ *e*(*s*_*t*_, *r*_*t*_; *ψ*_*t*_) depends on how incoming sensory stimuli are encoded in neural responses via the encoding parameters *ψ*_*t*_. In order to improve performance on this task, the system can adapt the encoding parameters over time. To do this, an observer uses the neural responses to construct a posterior belief *p*(*c*_*t*_|*r*_*τ*≤*t*_; *ψ*_*t*_) about the current context. This belief can then be used to construct a point estimate *ĉ*_*t*_ of the current context, and a point prediction of the future context. This prediction can then used to adapt the upstream encoding parameters via a feedback loop. Note that in practice, the current estimate provides a good approximation of this prediction in scenarios where the context changes with a small probability or by a small amount per unit time; we consider such scenarios for the remainder of the paper.

We evaluate the system’s overall performance by averaging the instantaneous task error over ongoing changes in context. If the system spends a total time *T*_*c*_ in each of *N*_*c*_ different contexts, this global (i.e., context-averaged) error can be written as:

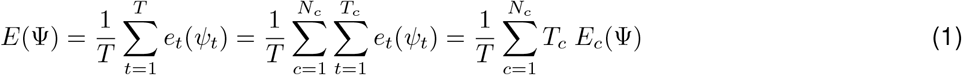

where 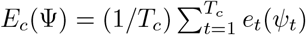 is a local (i.e., context-specific) error rate.

In order to achieve low global error, the encoding parameters *ψ*_*t*_ ∈ Ψ should be adapted to the statistics of incoming stimuli. In the simplest case, exemplified in Fig 1c, the system can use a fixed set of parameters Ψ ≡ *ψ*^global^ that are chosen to minimize the global error in Eq. 1, and that are thereby adapted to the set of all contexts:

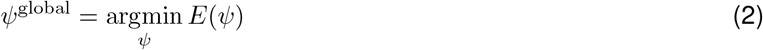

We refer to this as the “static” code.

The system can further improve its performance by adapting these parameters to each individual context. In this case, the system can maintain a set of parameters 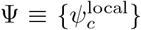 that are optimal for each context *c* by minimizing the local error rate *E*_*c*_. With perfect knowledge of the current context *c*_*t*_ at a given time *t*, an “oracle” code can then select among these context-optimal parameters, 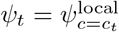. In the absence of perfect knowledge, the system can instead use a context *estimate ĉ*_*t*_ to select among these same parameters, 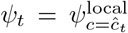; we refer to this as the “naively-adaptive” code. Note that a code is specified both by the method of optimizing the parameters (here, minimizing either global or local error, as indicated by the superscript), and the method of selecting among these parameters (using no context knowledge, perfect context knowledge, or an estimate of context, as indicated by the subscript). These properties are summarized in Fig 2c.

Since no real-world estimation strategy is perfect, any system that adapts based on a context estimate will face times when this estimate is correct and the system is matched to the current context (*ĉ*_*t*_ = *c*_*t*_), and times when this estimate is incorrect and the system is mismatched (*ĉ*_*t*_ ≠ *c*_*t*_). This observation allows us to decompose the global error in Eq. 1:

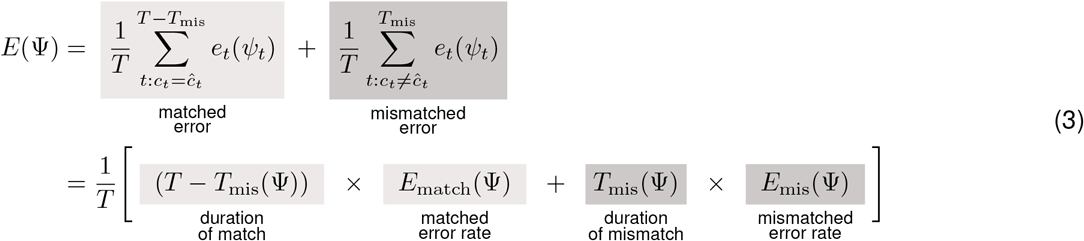

where (*T* − *T*_mis_) and *T*_mis_ denote the total duration of time that the system is “matched” or “mismatched”, respectively, and *E*_match_ and *E*_mis_ denote the respective error rates during these times. Periods of mismatch can lead to high errors if either the duration of these mismatches is long, or if error rates during periods of mismatch are high.

To date, theoretical studies of sensory coding have focused on the design of sensory codes that are optimally matched to the current context, and thus minimize the matched error rate *E*_match_ given perfect context knowledge [30, 35, 42–44]. Here, we extend these theories by observing that when the system does not have perfect knowledge of a context that is changing in time, its performance is determined by the *interplay* between the matched error rate and two additional factors: the mismatched error rate *E*_mis_, and the duration of the mismatch *T*_mis_. These three factors are intrinsically coupled and must therefore be balanced in order to achieve high performance.

The naively-adaptive code, which minimizes the matched error rate given the system’s current context estimate, is the most direct extension of previous work. This code, which prioritizes local task performance given a current context estimate, suffers from high task errors when this estimate is faulty. To reduce these errors, we construct codes that minimize either the duration of mismatch or the mismatched error rate (Fig 2c). The former prioritizes accurate inference of the underlying context (with parameters 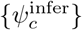 selected via a context estimate, 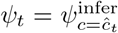); the latter prioritizes task performance in contexts other than the locally, but potentially incorrectly, estimated context (with parameters 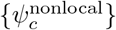 selected via a context estimate, 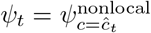). We interpolate between these and the naively-adaptive code by constructing families of so-called “biased” codes that achieve different balances between these priorities. We index these codes with a bias parameter *α*_B_ ∈ {*α*_I_, *α*_NL_} that takes a value of 0 in the naively-adaptive case that prioritizes local task performance, and a value of 1 in a fully-biased case that prioritizes either inference (with bias parameter *α*_I_), or nonlocal task performance (with bias parameter *α*_NL_).

Fig. 2d-e exemplifies how we dissect the relative contributions of different terms to the global error (note that we report the relative duration of mismatch *T*_mis_*/T*, which, when multiplied by *E*_mis_, gives the total mismatched error, as shown in Fig 2d). While biased codes are designed to reduce either the duration or error rate of mismatches, they are not guaranteed to reduce both simultaneously (Fig 2d), and therefore are not guaranteed to reduce the overall mismatched error. Moreover, biased codes can only increase matched error over the naively-adaptive code, which minimizes this error by construction (Fig 2e, left). However, there are settings in which biased codes significantly reduce mismatched error, thereby overcoming these increases in matched error and ultimately improving overall performance (Fig 2e, right). As we next show, the specific pattern of interplay between these different sources of error is scenario-specific, and thus different forms of biasing (i.e., toward either inference or nonlocal task performance) can lead to better or worse global performance under different conditions.

#### Performance tradeoffs and optimal coding schemes are scenario-dependent

To illustrate how global performance depends on the interaction between different sources of error, we consider a simple analytical approximation of the global error in Eq. 3. Each of the terms in this equation implicitly depends on the bias parameter *α*_B_ through the coding parameters *ψ*_*t*_ ∈ Ψ:

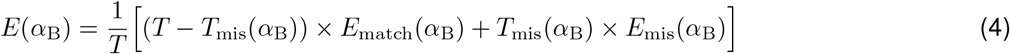

We can measure the properties of biased codes with respect to baseline properties of the naively-adaptive code, when *α*_B_ = 0. We denote these baseline properties with a superscript ‘0’, such that 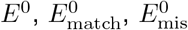, and 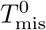 respectively denote the global error, matched error rate, mismatched error rate, and duration of mismatch of the naively-adaptive code (Fig 3a). The baseline properties 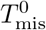 and 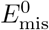 reflect the complexity of a dynamic sensory environment. In environments where context changes are difficult to infer, 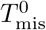 will be high; similarly, in environments where context mismatches are costly, 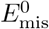 will be high. We can thus manipulate these baseline properties as a proxy for manipulating environmental complexity, and we can then analyze a particular code across a range of different environmental scenarios. Within any given environment (defined here by a specific combination of 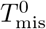 and 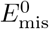), optimal performance will require a particular balance of the terms in Eq. 4. A family of biased codes can thus be defined in terms of the balance that it achieves.

**Figure 3:**
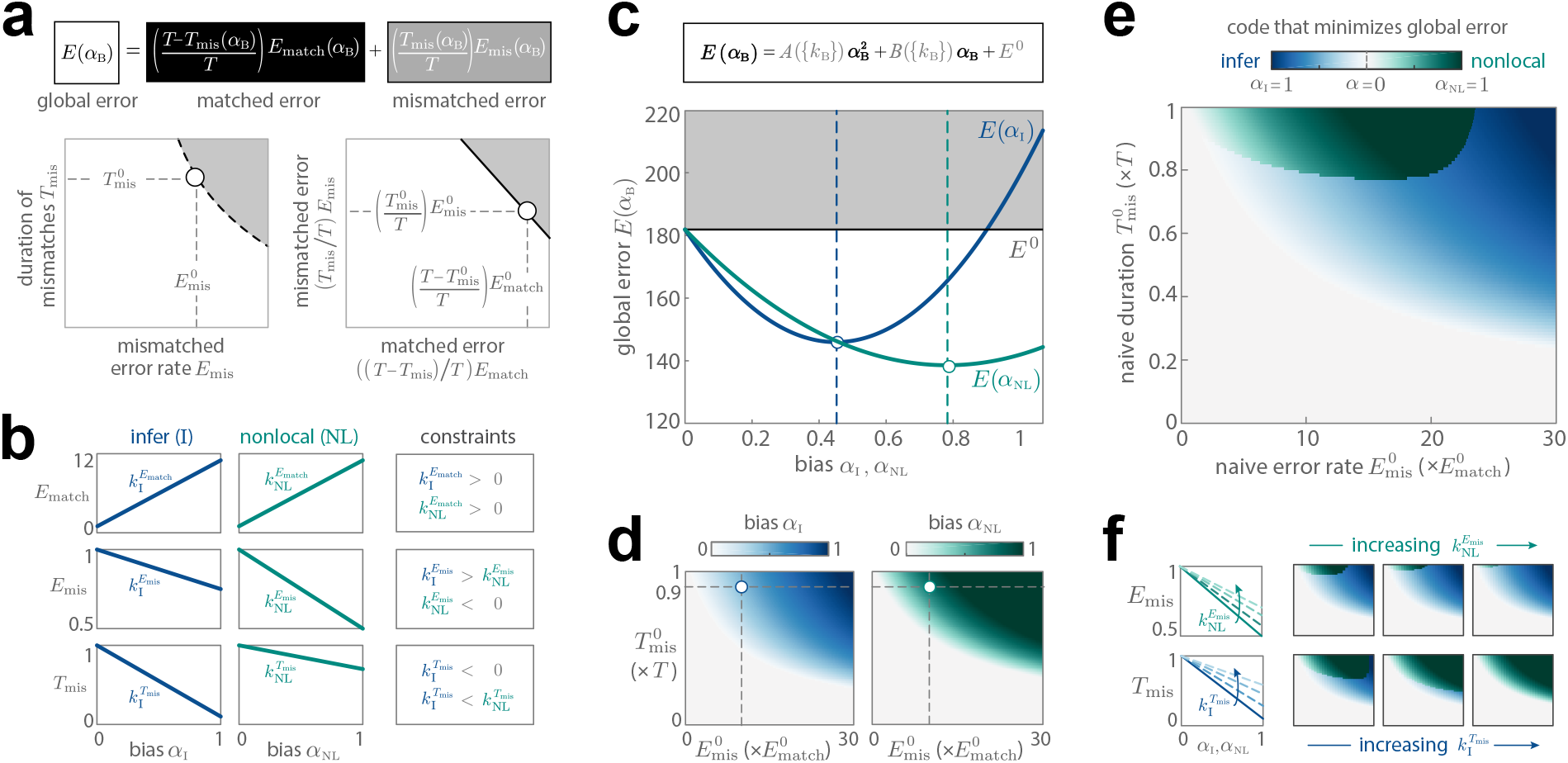
Optimal codes are scenario-dependent. **a)** Global error can be expressed as a function of the bias parameter *α*_B_, where B ∈ {I, NL} denotes whether codes are biased toward inference or nonlocal task performance, respectively. As in Fig 2d-e, we decompose error into matched and mismatched contributions (right panel), and further decompose mismatched error into the duration and error rate of mismatches (left panel). We measure changes in error relative to the naively-adaptive code (open markers; error terms indexed by superscript ‘0’). **b)** We approximate the matched error rate (*E*_match_, top row), mismatched error rate (*E*_mis_, middle row), and duration of mismatch (*T*_mis_, bottom row) as linear functions of the bias parameters *α*_I_ (left column) and *α*_NL_ (middle column). The nature of biasing imposes constraints on the set of linear coefficients {*k*_B_ } (right column; see Methods for details). Linear approximations are shown for *k*_I_ = 10, − 0.25, − 0.9 and *k*_NL_ = 10, − 0.5, − 0.3 (top, middle, bottom panels, respectively); the same values are used for the remaining panels. **c)** In this approximation, the global error depends quadratically on the bias parameter *α*_B_. Codes biased toward inference (*E*(*α*_I_); blue dashed line) and nonlocal task performance (*E*(*α*_NL_); green solid line) reduce global error below that of the naively-adaptive code (*E*^0^; gray line). The best inference code is achieved for *α*_I_ = 0.42 (blue dashed line); the best nonlocal code, and the code that results in the lowest global error, is achieved for *α*_NL_ = 0.73 (green dashed line). Shown for 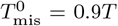 and 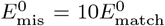, where *T* = 10^4^ and 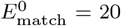. d) In different environments (parametrized by variations in 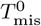 and 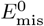), the best inference code and nonlocal code are achieved for different values of *α* _I_ (left panel, blue) and *α*_NL_ (right panel, green), respectively. Dotted lines mark the values used in (**c**). **e)** Depending on the environment, the optimal code that minimizes global error will either be biased toward inference (blue region) or toward nonlocal task performance (green region). The value of the bias parameter that achieves this minimum error can vary from 0 (i.e., a naively-adaptive code; white regions) to 1 (i.e., a fully biased code; full color saturation). **f)** Changing the balance of errors by modifying the coefficients {*k*_B_} will change the type and bias of the optimal code. Increasing 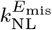 will reduce the extent to which biasing toward nonlocal task performance decreases the mismatched error rate *E*_mis_, and will thus reduce the domain for which this type of code is optimal (marked by the decreasing size of green regions; upper row). Shown for 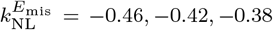 (left, middle, right panels, respectively). Analogously, increasing 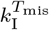 will reduce the extent to which biasing toward inference decreases the duration of mismatches *T*_mis_, and will thus reduce the domain for which this type of code is optimal (marked by the decreasing size of blue regions; lower row). Shown for 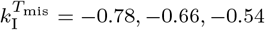 (left, middle, right panels, respectively). Axes of heatmaps span the same ranges as in (**d**,**e**).

By construction, the naively-adaptive code minimizes the matched error rate; codes that bias toward inference (by increasing *α*_I_) or nonlocal task performance (by increasing *α*_NL_) will monotonically increase this error. In doing so, the former family of codes will monotonically reduce the duration of mismatch, *T*_mis_(*α*_I_), and the latter family will monotonically reduce the mismatched error rate, *E*_mis_(*α*_NL_). We can therefore use linear functions to approximate the dependence of these terms on *α*_B_ (Fig 3b, left and middle columns):

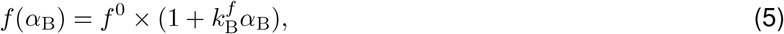

where *f* ∈ {*E*_match_, *E*_mis_, *T*_mis_} denotes different sources of error, and the coefficients 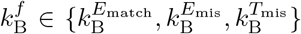 denote the corresponding slopes of their linear dependence on *α*_B_. When written in this way, these coefficients specify the proportion by which the properties of a fully-biased code (with *α*_B_ = 1) will deviate from the naively-adaptive code (with *α*_B_ = 0), subject to constraints imposed by the particular form of biasing (Fig 3b, right column; see Methods for details).

Given this linear approximation, we can now express the global error as a quadratic function of *α*_B_:

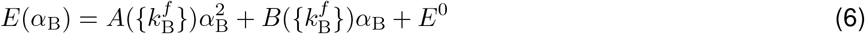

where *A*(·) and *B*(·) each depend on the set of linear coefficients 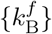 (see Methods for full derivation). By varying these coefficients, one can identify the type of code (naively-adaptive, biased toward inference, or biased toward nonlocal task performance), the degree of bias (specified by *α*_B_), and the balance of errors (specified by 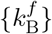) that minimizes global error for a particular environment.

To illustrate this interplay, we consider an example set of linear coefficients that specify the balance of errors achieved by a family of codes that bias toward inference (with 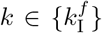), and a second family of codes that bias toward nonlocal task performance (with 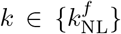) (Fig 3b). We begin by evaluating the performance of these codes in a single environment. Both families of codes reduce the global error below that of the naively-adaptive code (Fig 3c), but the latter family of codes achieves the lowest global error with a bias of *α*_NL_ = 0.73.

If we now take these same families of codes and evaluate them in different environments, we see that different values of *α*_I_ and *α*_NL_ are required to minimize global error within each family (Fig 3d). In environments where context changes are easy to detect and the cost of mismatches is low (Fig 3d, white regions in bottom left corners of each panel), biasing does not decrease the global error, and the naively-adaptive code is optimal. The more complex the environment, however, the stronger the bias (and thus the larger the value of *α*_B_) that is required to minimize the global error. Moreover, depending on the properties of the environment, the family of codes that achieves the lowest global error might change from the family biased toward nonlocal task performance (Fig 3e, green region) to the family biased toward inference (Fig 3e, blue region). The extent of these regimes is controlled by the specific balance of errors achieved by each family of codes (Fig 3f).

This simple approximation highlights the multiple factors that shape global performance. As we will show next, the explicit dependence on *α*_I_ and *α*_NL_ can be complex and itself scenario-dependent, and the resulting balance of errors can be shaped by factors such as number and diversity of statistical contexts, the measure of task error, and the nature of coding constraints. The resulting performance will nevertheless be shaped by this balance of errors, and the code that optimizes performance will vary depending on the scenario. For the remainder of the paper, we focus on those scenarios in which biased codes reduce global error and thereby improve global performance over the naively-adaptive code. We measure this improvement relative to the difference in global error between the static and oracle codes, and we demonstrate that this improvement holds across a range of different scenarios, and even for simple approximative coding schemes. However, we note that global error is not the only coding objective for assessing performance, and this framework enables a rigorous analysis of a range of different objectives depending on the scenario and task at hand. For example, some scenarios might necessitate rapid detection of changes in the environment, and might specifically require codes that minimize the time of adaptation, *T*_mis_. Other scenarios might necessitate maintaining instantaneous error levels below a certain value, and might specifically require codes that minimize *E*_mis_. Yet other codes might prioritize accuracy when the environment is stable and might minimize *E*_match_. By understanding how these different terms contribute to the global error in Eq. 3, we can dissect the properties of adaptive dynamics that reflect a range of different coding objectives, something that we will revisit in comparisons to data.

#### Locally suboptimal performance can improve global performance

We now revisit our linear-nonlinear Poisson neuron (Fig 1) through the lens of this framework (Fig 4). Here, we explicitly model the temporal dynamics of the environment, which we take to switch between each of the nine different contexts at a small but fixed probability per time.

**Figure 4:**
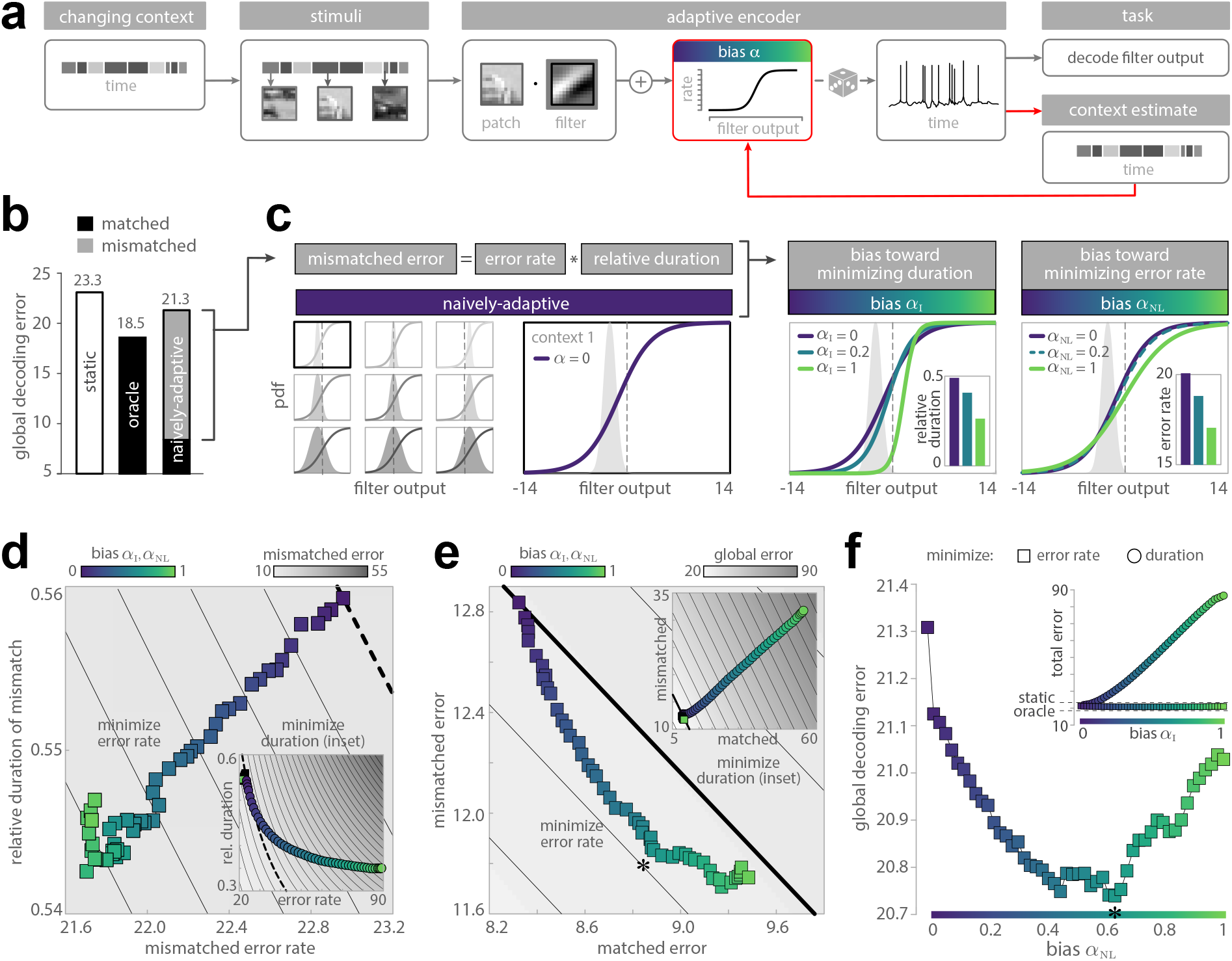
Biased codes improve global performance. **a)** We revisit the example Poisson neuron shown in Fig 1 through the lens of the framework shown in Fig 2. Here, the environment can switch between each of the nine contexts with a small but fixed probability per time. Rather than having perfect knowledge of this context, the neuron must construct an estimate of context that can be fed upstream and used to adapt the encoding nonlinearity (red arrow) for the task of decoding filter outputs. **b)** A naively-adaptive code that adapts based on a context estimate achieves performance intermediate between static and oracle codes. A large fraction of the global error arises during periods of mismatch (gray), when the context estimate is inaccurate. **c)** The naively-adaptive code uses its context estimate to select among the context-optimal nonlinearities shown in Fig 1b and reproduced here. To reduce mismatched error, we bias these nonlinearities toward inference (middle panel) or nonlocal task performance (right panel). The former is achieved by biasing each nonlinearity towards the maximally discriminative nonlinearity, which reduces the duration of mismatch (inset). The latter is achieved by biasing each nonlinearity towards the nonlinearity that accurately decodes stimuli in other contexts, which reduces the mismatched error rate (inset). **d)** Codes biased toward inference (circles) result in relatively large mismatched error rates (inset). Codes biased toward nonlocal task performance (squares) show a reduction in both the duration and error rate of mismatches, and thereby achieve lower mismatched error (main panel). Solid lines mark contours of constant mismatched error. **e)** Codes biased toward inference (circles) increase both matched and mismatched error (inset). Codes biased toward nonlocal task performance (squares) exhibit a tradeoff between matched and mismatched error (main panel). The code that optimally balances matched and mismatched error (and thereby achieves the lowest global error) is marked by a star. Solid lines mark contours of constant global error. **f)** As bias increases, codes biased toward inference (circles) quickly exceed the global error produced by the static code, and continue to monotonically increase thereafter (inset). In contrast, codes biased toward nonlocal task performance (squares) achieve total errors that are intermediate between oracle and static codes, and are minimized for an intermediate bias of *α*_NL_ = 0.61. For display purposes, all error values in panels d-f, except for *α*_NL_ = *α*_I_ = 0, were smoothed with a moving window of width 8.

As described earlier, these contexts specify the distributions of outputs that are produced by the neuron’s fixed linear filter across an ensemble of natural scenes; we approximate these as Gaussian distributions with different means and variances (i.e., *p*(*s*_*t*_|*c*_*t*_) = 𝒩 (*s*_*t*_; *µ*(*c*_*t*_), *σ*(*c*_*t*_)^2^)). At a given time *t*, a filter output *s*_*t*_ is sampled from the distribution specified by the current context *c*_*t*_ and transformed into a Poisson-distributed spike count *r*_*t*_ via a sigmoidal nonlinearity with adaptable slope *k* and offset *s*_0_. The finite dynamic range of this nonlinearity controls the maximum firing rate of the neuron, and thus limits the precision with which incoming stimuli can be encoded in the response *r*_*t*_. Neural responses over a window of time Δ*T* are then used to update a maximum-likelihood estimate *ĉ*_*t*_ of the current context, which can then be used to adapt the encoding parameters *ψ* = {*k, s*_0_ } for the task of accurately decoding the filter output. We measure the decoding error at time *t* as the squared error between the maximum-likelihood estimate of the filter output, *ŝ*_*t*_, and its true value *s*_*t*_ (i.e., *e*_*t*_ = (*s*_*t*_ − *ŝ*_*t*_)^2^). We then average this error over time, and thus over many context switches, to compute the global decoding error.

We first construct a naively-adaptive code that uses its context estimate to select among the local, context-optimal nonlinearities shown in Fig 1c. The parameters 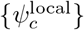 of these nonlinearities were chosen to minimize the error in decoding filter outputs sampled from each individual context *c*. We compare the performance of this code to the performance of an oracle code, which uses the true context to select among these same nonlinearities (black bar, Figs 1d and 4b), and a static code, which uses a single global nonlinearity whose parameters *ψ*^global^ were chosen to minimize the error in decoding filter outputs sampled from all contexts (white bar, Figs 1d and 4b). The global decoding error produced by the naively-adaptive code is intermediate between the errors produced by the static and oracle codes, and has a large contribution from periods of mismatch (Fig 4b).

To reduce the impact of these periods of mismatch, we bias this naively-adaptive code (Fig 4c, left) toward inference (Fig 4c middle, with parameters 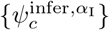), or toward nonlocal task performance (Fig 4c right, with parameters 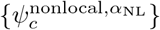). In the former case, we bias the parameters of the nonlinearity optimized for a particular context *c* toward the nonlinearity that best discriminates between contexts (see Methods). In the latter case, we bias the same parameters toward the nonlinearity that minimizes the error in decoding filter outputs sampled from all other contexts *c*′ ≠ c; we sample these filter outputs in proportion to the probability of estimating the true context *c*′ as the erroneous context *c* (i.e., 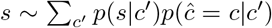. In both cases, we do this by constructing a weighted combination of the naively-adaptive and fully-biased parameters, where *α*_I_ and *α*_NL_ specify the weight between these two sets of parameters; note that this approach is an approximation to direct optimization that is computationally tractable and allows us to illustrate the existence of different coding regimes. Although these two different forms of biasing lead to seemingly subtle differences in the resulting nonlinearities, the impact on overall performance is substantially different (Fig 4d-f). Biasing toward nonlocal task performance by reducing the mismatched error rate also leads to a weak reduction in the duration of the mismatch, and thereby a reduction in the total mismatched error (Fig 4d, square markers). This reduction in mismatched error is accompanied by an increase in matched error, and the global error (the sum of these two terms) is thus minimized for an intermediate bias of *α*_NL_ = 0.61 (starred points in Fig 4e-f). In comparison, biasing toward inference by reducing the duration of mismatch leads to a significant increase in the mismatched error rate (Fig 4d; inset). This leads to higher mismatched error, which, together with the dramatic increase in the matched error (Fig 4e; inset), ultimately results in higher global error (Fig 4f; inset).

Together, these results highlight that biasing away from the strategy that would be locally optimal if the context were stationary can improve global performance when the context is changing. The finding that global error is reduced by biasing toward a reduction in the error rate, but not the duration, of mismatches could be due in part to the suboptimality of our context inference strategy. Optimal context inference would require an ideal Bayesian observer model that could leverage the uncertain and continually-changing estimate of the current context, which would in turn improve the accuracy of inferring context switches. With such an ideal observer, it is possible that biasing toward a reduction in the duration of mismatches could improve global performance, rather than hinder it. In what follows, we examine a scenario in which the ideal Bayesian observer can be easily derived and implemented by a simple neural encoder.

### A normative explanation of the dynamics underlying variance and mean adaptation

We now consider a simplification of the previous scenario that mimics one of the most widely-studied adaptation paradigms in sensory neuroscience—adaptation to changes in stimulus variance and mean [5, 10, 11, 16, 23, 24, 45]. In this scenario, where stimulus statistics are highly controlled, the ideal observer model for inferring changes in these statistics is well known and provides a continuous estimate of the changing sensory context [31]. We construct a family of sensory codes that use this estimate to optimally balance different sources of error (Fig 5), and we show that these codes explain a diversity of adaptive dynamics that have been observed experimentally (Fig 6). The simplicity of this setting, and its strong theoretical and experimental grounding, enables us to dissect the dynamics that support optimal adaptation and compare these dynamics to experimental data.

**Figure 5:**
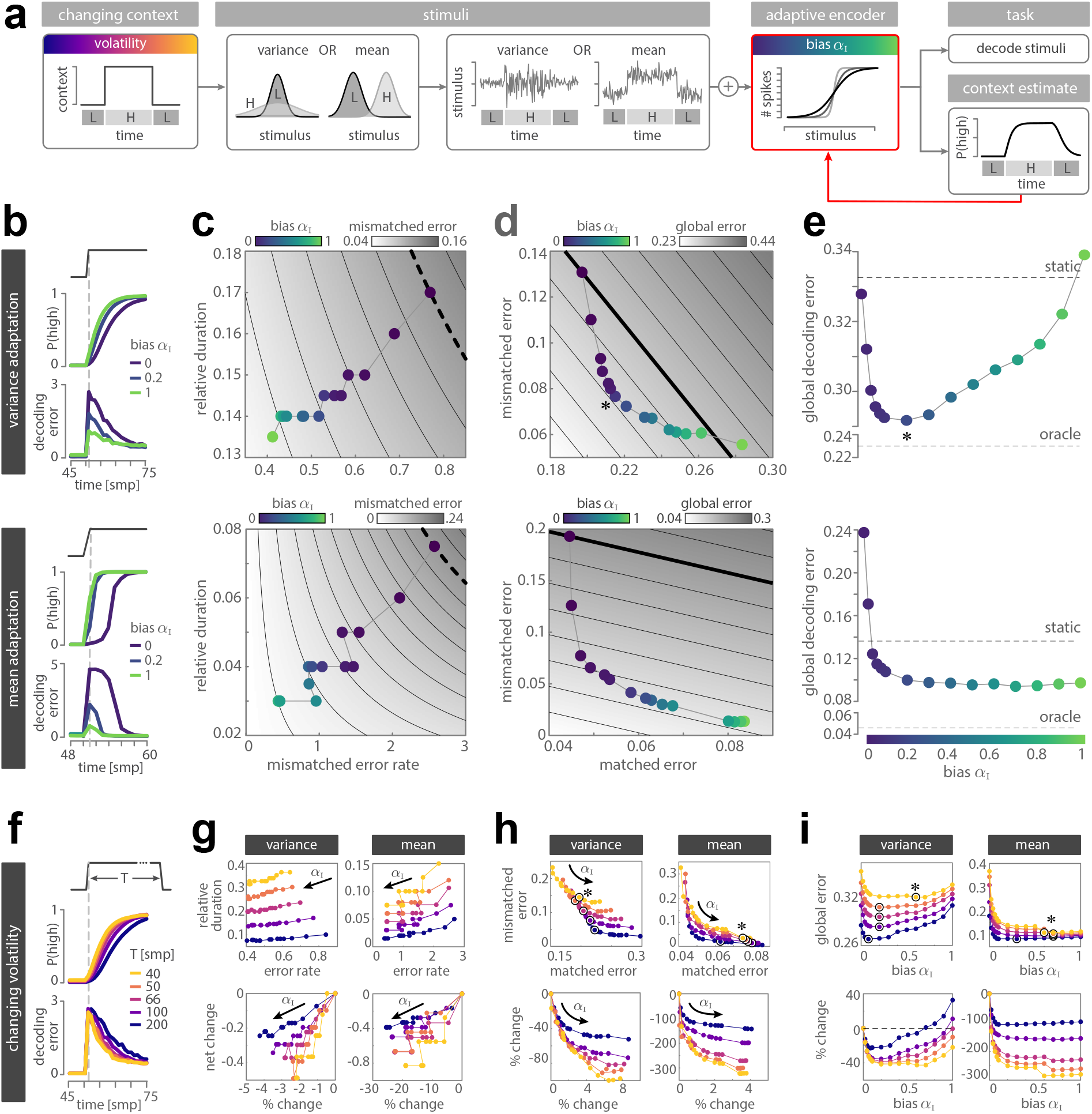
Biased codes improve performance across a range of different stimulus conditions. **a)** We consider a simple model neuron in an environment that can switch between a low (L) and high (H) context over time. This context signals either the mean or variance of a Gaussian stimulus distribution. The neuron encodes incoming stimuli in a discrete response via a saturating nonlinearity. This response is used to estimate the posterior probability 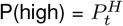 that the environment is in the high context, which can be fed upstream (red arrow) used to adapt the encoding nonlinearity for the task of accurately decoding stimuli. We consider a family of codes that bias toward maintaining an accurate estimate of context (blue-green colormap), and we track the performance of these codes in settings with varying volatility (purple-yellow colormap). **b)** Following an abrupt increase in variance (upper panel) or mean (lower panel), the system updates its posterior belief P(high) that the environment is in the high context. This is accompanied by a transient increase in decoding error. **c)** Biasing the code toward inference leads to shorter periods of mismatch and lower mismatched error rates during both variance adaptation (upper panel) and mean adaptation (lower panel). Solid lines indicate contours of constant mismatched error. **d)** Biased codes exhibit a tradeoff between matched and mismatched error during both variance adaptation (upper panel) and mean adaptation (lower panel). Solid lines indicate contours of constant global error, and the code that achieves lowest global error is marked with a star. **e)** During variance adaptation (upper panel), most biased codes outperform the static code, and optimal performance is achieved for an intermediate bias of *α*_I_ = 0.2 (star). During mean adaptation (lower panel), the static code and all biased codes outperform the naively-adaptive code (*α*_I_ = 0). Larger biases improve performance. **f)** As volatility increases, the posterior is faster to update and the error faster to decay (shown for the naively-adaptive code in response to an increase in variance). **g-i)** Biased codes produce qualitatively consistent patterns of performance regardless of volatility (upper panels), but the relative impact of biasing the code increases as volatility increases (lower panels). Higher volatility leads to relatively longer periods of mismatch (**g**, upper), higher mismatched errors and lower matched errors (**h**, upper), and higher global errors (**i**, upper). The percent change in each of these errors (measured relative to the naively-adaptive code and scaled by the difference in error between the static and oracle codes) increases as volatility increases (lower row), and the global error is thus minimized for larger values of bias (**h-i**; circled markers in upper panels).

**Figure 6:**
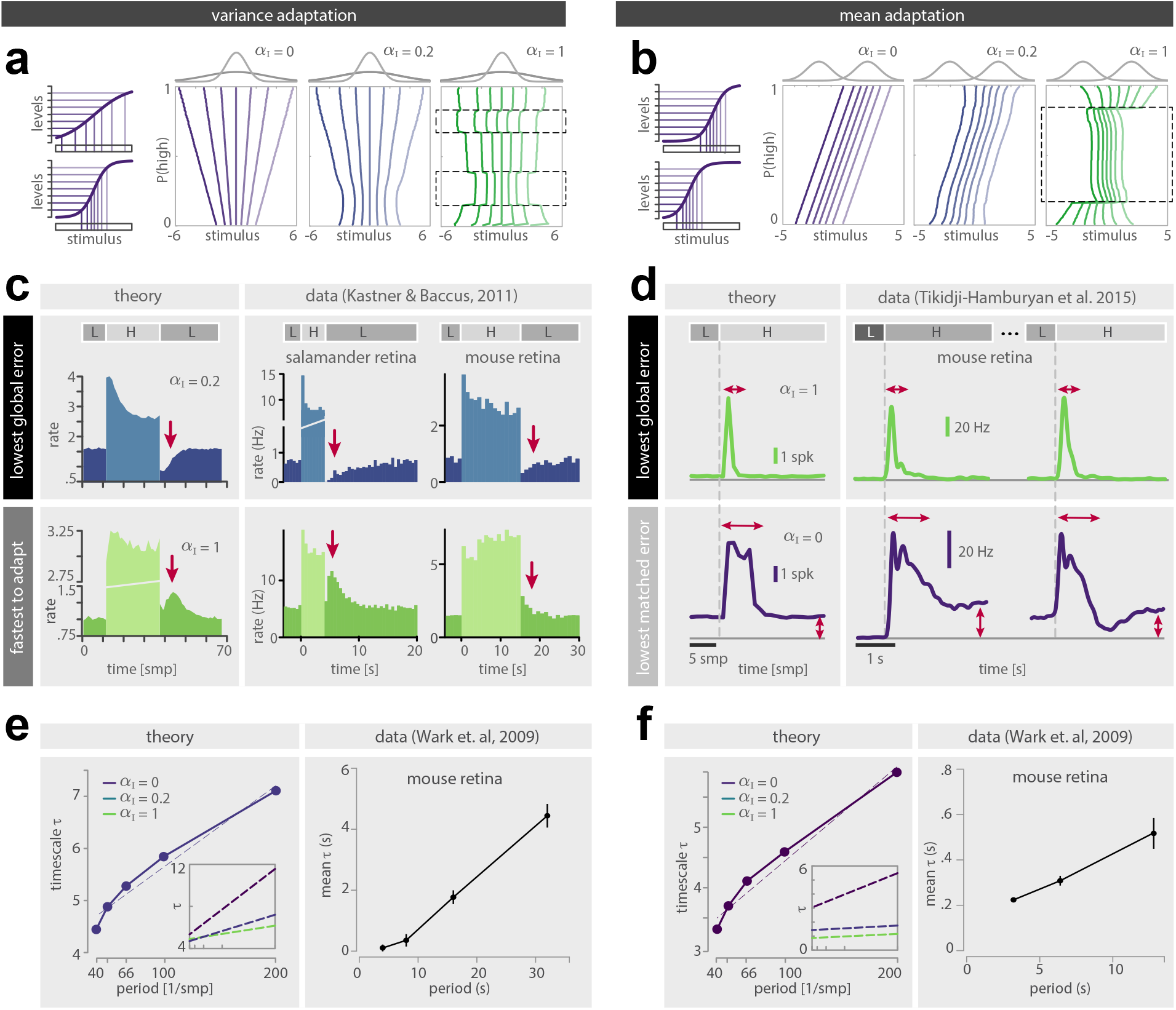
Biased codes capture diverse profiles and timescales of adaptive dynamics. **a, b)** Optimal nonlinearities vary as a function of the system’s posterior belief 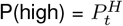 about the current context, and they differ between the naively-adaptive (purple; left), fully-biased codes (green; right), and intermediate (blue; center) codes. To visualize these changes, we use vertical lines to differentiate sets of stimuli that are encoded in different response levels (small left schematics in **a** and **b**; exemplified for the naively-adaptive code for P(high)=1 (upper) and P(high)=0 (lower)). Regions where these lines are closer together mark steeper portions of the nonlinearity. **a)** When the naively-adaptive code (left) is certain of either the high- or low-variance context (P(high) near 1 or 0), the nonlinearity is either broad or narrow, respectively, and linearly scales between these extremes. When the fully-biased code (right) is more certain of the high-variance context (P(high) above 0.5 and below 1), the nonlinearity broadens and shifts toward the tail of the high-variance stimulus distribution (upper dashed box); when the code is more certain of the low-variance context (P(high) above 0 and below 0.5), the nonlinearity sharpens and centers (lower dashed box). The optimal code (center) shares features of the naively-adaptive and fully-biased codes. **b)** When the naively-adaptive code (left) is certain of either the high- or low-mean context (P(high) near 1 or 0), the nonlinearity is centered under the high- and low-mean distributions, respectively, and linearly scales between these extremes. When the fully-biased code (right) is certain of either context, the nonlinearity is broad shifted toward the distribution that signals the other context; when the code is uncertain, the nonlinearity is sharp and centered between the two distributions (dashed box). An intermediate code (center) shares features of the naively-adaptive and fully-biased codes. **c, d)** Average firing rates change in response to a step increase and decrease in variance (**c**) and mean (**d**). **c)** The code that minimizes global error (upper left; *α*_I_ = 0.2) produces a transient drop in firing rate following a change from high (H) to low (L) variance (red arrows); in contrast, the code that maximizes adaptation speed (lower left; *α*_I_ = 1) produces a transient increase in firing rate. Similar dynamics have been observed in “adapting” (upper row) and “sensitizing” (lower row) ganglion cells in the retina of salamander (middle column) and mouse (right column) in response to step changes in contrast (figure courtesy of David Kastner; [16]). **d)** The code that minimizes global error and simultaneously maximizes adaptation speed (upper left; *α*_I_ = 1) produces a sharp and transient increase in firing rate following an change from low (L) to high (H) mean (red arrows). In contrast, the code that minimizes the matched error rate (lower left; *α*_I_ = 0) produces a higher baseline firing rate and a longer and doubly-peaked transient increases following a change in mean (lower left; *α*_I_ = 1). Similar dynamics have been observed in the mouse retina, shown for two ON ganglion cells (different rows) responding to a step increase in luminance at lower (middle column) and higher (right column) light levels (figure courtesy of Thomas Münch; [46]). **e, f)** The timescale of adaptation to changes in variance (**e**, left) or mean (**f**, left) increases with the period of these changes. This relationship is well approximated by a linear fit (dashed line) whose slope and offset varies as a function of the bias *α* (inset). In mouse retinal ganglion cells (**e, f**, right), the timescale of adaptation exhibits similar scaling with the switching period of the stimulus distribution (figure reproduced from [45]). Timescales were estimated using an exponential fit; see Methods.

#### Performance tradeoffs generalize across variations in stimulus statistics

To model this widely-studied paradigm, we consider a simplification of the setup in Fig 4a, shown in Fig 5a, in which the environment switches between one of two contexts—a ‘high” context (*c* = *c*_*H*_), and a “low” context (*c* = *c*_*L*_)—at a small but fixed probability per time. We separately vary the volatility of these context switches by adjusting this probability. We take this latent context to specify either the variance or the mean of a Gaussian stimulus distribution (i.e., 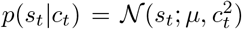) or 𝒩 (*s*_*t*_; *c*_*t*_, *σ*^2^), respectively). This corresponds to two scenarios in which either the variance of incoming stimuli is changing over time (“variance adaptation”), or the mean is changing over time (“mean adaptation”). Stimuli from these distributions are then encoded by a simple model neuron. Specifically, at a given time *t*, a stimulus *s*_*t*_ is drawn from the distribution specified by the current context *c*_*t*_, transformed via a sigmoidal nonlinearity with adaptable slope *k* and offset *s*_0_, distorted by noise, and finally discretized into one of *n* different response levels to produce a response *r*_*t*_. These response levels can be interpreted as spike counts or discriminable firing rates, and their discrete nature limits the precision with which incoming stimuli can be encoded in the response *r*_*t*_. This response is then linearly decoded to extract a stimulus estimate *ŝ*_*t*_, which is used to update an internal estimate of the current context *ĉ*_*t*_. To construct this estimate, we use a optimal Bayesian observer that is identical in form to the observer derived in [31], but uses decoded stimulus estimates rather than raw stimulus values to update its internal context estimate. This observer first computes the posterior probability 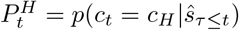 that the environment is in the high context at time *t*, and then averages this posterior over contexts to produce a continuous context estimate *ĉ*_*t*_. This context estimate is then used to adapt the encoding parameters *ψ* = {*k, s*_0_ } for the task of accurately decoding the stimulus. We measure the decoding error at time *t* as the squared error between the decoded stimulus estimate *ŝ*_*t*_ and raw stimulus value *s*_*t*_ (i.e., *e*_*t*_ = (*s*_*t*_ − *ŝ*_*t*_)^2^). We then average this error over time and context switches to compute the global decoding error.

We first construct a naively-adaptive code whose encoding parameters are chosen to accurately decode stimuli given the system’s current context estimate. Note that because this estimate is continuous, we construct a set of encoding parameters that can take on a continuum of values. To this end, the parameters 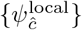 are chosen to minimize the error in decoding stimuli sampled from the system’s estimate of the stimulus distribution, *p*(*s*_*t*_|*ĉ*_*t*_). We then bias this code toward inference by determining the encoding parameters 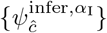 that minimize a weighted combination of the error in decoding stimuli and the error in estimating the context, where *α*_I_ specifies the weight between these two error terms. We empirically found that this strategy, rather than one biased toward nonlocal task performance, produced the largest reduction in global error, and we therefore focus on this strategy for the remainder of this section. We compare the performance of these biased codes to a static code whose parameters *ψ*^global^ are chosen to minimize the average error in decoding stimuli sampled from both contexts, and an oracle code whose parameters 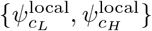 are chosen to minimize the average error in decoding stimuli sampled from either the low or high contexts, respectively.

Fig 5b shows the dynamic evolution of the posterior belief and the decoding error in response to a step increase in stimulus variance (upper panel) or mean (lower panel). In both cases, the naively-adaptive code is the slowest to respond to this step increase, and produces the largest and longest transient errors following this increase. We partition these dynamics into times when the system’s posterior belief is within or outside a fixed tolerance of the true context (corresponding to times when the context estimate is matched or mismatched, respectively), and we analyze the contributions of these dynamics to global error (Fig 5c-e). For both variance and mean adaptation, biased codes reduce both the duration and error rate of mismatches as compared to the naively-adaptive code (Fig 5c), consistent with the dynamics shown in Fig 5b. This leads to an overall reduction in mismatched error but an increase in matched error, and thus the global error is minimized for a nonzero bias (starred points in Fig 5d-e). In the case of variance adaptation, all but the fully-biased code achieve lower global error than the static code, and optimal performance is achieved for an intermediate bias of *α*_I_ = 0.2. In the case of mean adaptation, the static code achieves lower global error than the naively-adaptive code. Biased codes reduce this error even further, substantially outperforming the naively-adaptive code. Together, these results are qualitatively consistent with those observed in Fig 4; however, note that here, the improvement in performance is achieved by codes that are biased toward reducing the duration, rather than the error rate, of mismatches.

We observe qualitatively similar results as we vary the volatility of context switches, regardless of whether these switches signal changes in variance or mean (Fig 5f-i). Higher volatility (i.e., more frequent context switches) leads to faster adaptation (Fig 5g), but ultimately results in relatively longer periods of mismatch (upper row of Fig 5g), larger mismatched errors (upper row of Fig 5h), and larger global errors (upper row of Fig 5i). Biasing, in turn, has a stronger relative impact on performance when the volatility of the environment is higher (lower row of Fig 5g-i). As a result, the more volatile the environment, the larger the bias that minimizes global error (Fig 5i).

#### Balancing matched and mismatched errors explains a diversity of adaptive neural dynamics

The improvements in performance shown in Fig 5 arise as a consequence of dynamically adapting the encoding nonlinearity of our model neuron, which in turn determines how the neuron’s limited response levels are allocated to different regions of the stimulus space. We can thus understand the performance of different codes in terms of how they allocate these response levels given the posterior belief about the underlying context (Fig 6a-b).

The naively-adaptive code uses a strategy that directly tracks the system’s posterior belief about the underlying context. When this context determines the variance of incoming stimuli, the system scales the width of the nonlinearity to track the belief, but it does not shift the location of the nonlinearity along the stimulus axis (Fig 6a, left). When the context determines the stimulus mean, the system shifts the location of the nonlinearity to track the belief, but it does not scale the width (Fig 6b, left).

In contrast, fully-biased codes implement a form of uncertainty-dependent change detection (Fig 6a-b, right). When the system’s posterior belief is close to 1 or 0 (i.e., the system is certain that the context is either high or low, respectively), the system shifts the nonlinearity away from the current context, thereby hedging its bets toward the change that is likely to occur. When the posterior belief approaches 0.5 and the system becomes uncertain, the system uses a nonlinearity that can better resolve this uncertainty. This manifests in different behavior depending on whether the stimulus variance or mean is changing in time. In the former case, increases in variance are easier to detect than decreases in variance [31], and so the behavior of the system is asymmetric to variations in uncertainty (Fig 6a, right). When the system is more certain that the variance is high, it uses a broad and shifted nonlinearity to resolve those stimuli of large magnitude that would signal high variances (upper dashed box in Fig 6a, right; note that we observe two symmetric solutions that are shifted toward either tail of the high variance distribution; the solution shown here produces fewer spike counts on average). On the other hand, when the system is more certain that the variance is low, it uses a sharp and centered nonlinearity to resolve stimuli of small magnitude that would signal low variances (lower dashed box in Fig 6a, right). In the latter case when the mean is changing, the behavior is symmetric; when the system is somewhat uncertain (regardless of whether it is more certain that the mean is either high or low), it uses a narrow nonlinearity that is centered between the two stimulus distributions, thereby enabling the system to accurately resolve stimuli that would differentiate distributions with high versus low means (dashed box in Fig 6b, right).

The nonlinearities in Fig 6a-b show the optimal partitioning of the stimulus space as a function of the system’s belief about the current context. In a dynamic setting, both this belief and the nonlinearities derived from it will be continually changing, which will together determine the firing rate of our model neuron. To extract this firing rate, we use an entropy coding scheme to recode the output of these nonlinearities. This parameter-free scheme assigns higher spike counts to response levels that are predicted to be used with lower frequency based on the system’s current belief about the underlying context (see Methods; SI Fig S1). The resulting firing rates are lower on average, and exhibit a range of different dynamics that vary depending on the bias *α*_I_ in a manner that resembles the diversity of neural dynamics observed experimentally in salamander, mouse, rabbit, and primate retina (Fig 6c-d; [16, 46– 48]).

For both mean and variance adaptation, codes that minimize global error produce firing rate dynamics that are qualitatively consistent with neural data (Fig 6c-d, upper row). In the case of variance adaptation, global error is minimized for *α*_I_ = 0.2, and the corresponding code exhibits firing rate dynamics that resemble the activity of so-called “adapting” cells [16, 47, 48]. When stimulus variance increases, these cells show an increase in firing rate followed by decay; when the stimulus variance decreases, the firing rate shows an abrupt drop followed by a recovery (Fig 6c, upper row). In the case of mean adaptation, global error is minimized for a range of *α*_I_ values between 0.2 and 1 (Fig 5e, lower panel); a value of *α*_I_ = 1 simultaneously minimizes the duration of mismatches, and is thus also the fastest to adapt. The corresponding code produces a low baseline activity and a sharp and transient increase in firing rate following a change in stimulus mean, resembling the activity of cells in mouse retina [46] (Fig 6d, upper row).

We additionally find cell classes whose whose dynamics resemble those of codes optimized for other terms in the error decomposition in Eq. 3, beyond global error. One class of so-called “sensitizing” cells show similar dynamics to adapting cells in response to an increase in stimulus variance, but they show qualitatively different dynamics following a decrease in stimulus variance [16, 47, 48]. These dynamics resemble those of codes that maximize the speed of variance adaptation, with *α*_I_ = 1 (Fig 6c, lower row). Another class of cells exhibits higher baseline activity and a prolonged, double-peaked response following an increase in stimulus mean [46]. These dynamics resemble those of codes that minimize matched error during mean adaptation, with *α*_I_ = 0 (Fig 6d, lower row). Thus, by varying a single parameter *α*_I_, this model generates a rich family of neural dynamics that closely resemble a diversity of empirical phenomena observed in different functional cell types. Importantly, each of these phenomena has a meaningful interpretation within our framework; they correspond to minimizing distinct error terms that arise when coding in dynamics environments. This suggests that different functional cell types are optimized to balance these different sources of error in order to maintain high performance in the face of changing stimulus statistics.

In all cases, the speed of adaptation depends on the rate of context switches. When this rate is low, the system expects the environment to be stable for longer periods of time, and thus requires more stimulus samples to be convinced that the environment has switched contexts. This leads to slower changes in the posterior belief (Fig 5f; SI Fig S2), and ultimately longer timescales of adaptation (Fig 6e-f, left panels). We observe this effect regardless of whether the system is adapting to changes in variance or mean, and regardless of the extent to which we bias the code. However, we find that the bias does affect the slope of this relationship; larger biases lead to shallower slopes and lower overall timescales (insets of Fig 6e-f). Similar relationships between the timescale of adaptation and the switching period of the stimulus have been observed in the fly, mouse, rat, and electric fish (Fig 6e-f, right panels; [10, 45, 49, 50]).

### Tradeoffs in receptive field adaptation to higher-order stimulus statistics

The previous sections illustrate the adaptive benefits of biasing away from locally optimal strategies in order to improve global performance. To illustrate the generality of these benefits beyond single neurons and low-order stimulus statistics, we consider a simple feed-forward network model of two visual neurons that use an adaptive set of linear receptive fields to jointly encode local patches of natural images (Fig 7a). This model falls within a broader class of linear neural models that have been used to both capture coding properties of real neurons [51] and predict coding properties from theoretical principles [52–54]. In order to maximize information about incoming images patches, these receptive fields must be adapted to their local statistical structure [52]. However, the statistics of these local image patches depend strongly on their location within a visual scene [55, 56]. Here, we use unsupervised learning to extract four statistical contexts that roughly correspond to the dominant cardinal orientation of a set of patches, and delineate different spatial regions of the scene (Fig 7b; see Methods).

**Figure 7:**
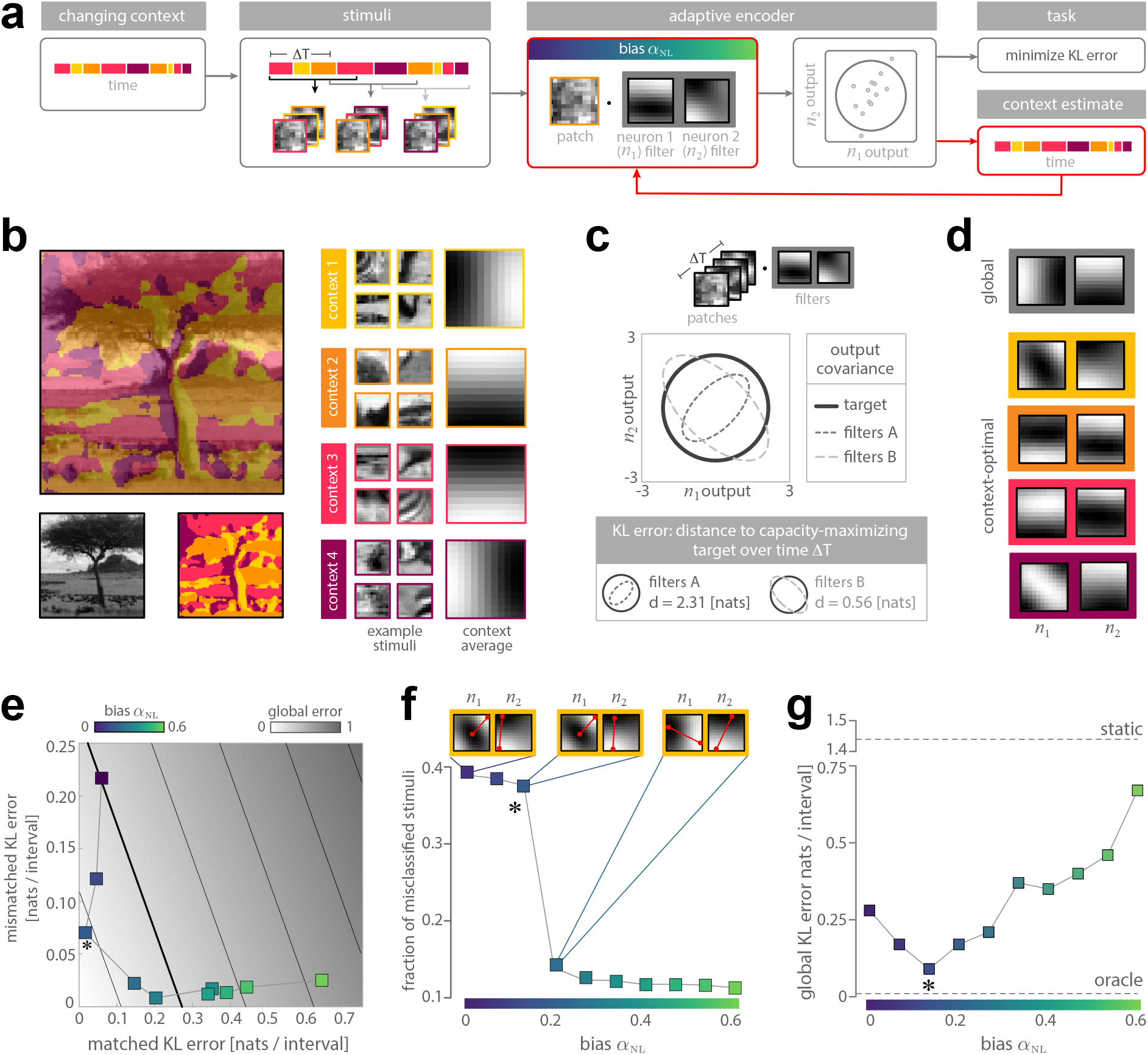
Feed-forward network can increase information transmission by biasing away from local stimulus statistics. **a)** A simple feed-forward network of two visual neurons uses linear receptive fields to encode incoming image patches. These patches originate from different contexts that can switch over time. The joint response of these neurons is used to construct an estimate of the current context, which then is fed upstream (red arrow) and used to adapt the receptive fields for the task of maximizing information transmission. **b)** Contexts with similar statistical structure correspond to different regions of a visual scene (left column). We consider four contexts that were discovered by unsupervised learning and that roughly correspond to the orientation of features within individual image patches (middle column; see Methods); the average image patch within each context summarizes the differences between contexts (right column). **c)** To quantify the error in information transmission during a time interval Δ*T* (schematic in upper row), we estimate the symmetrized KL-divergence (“KL error”) between the approximate distribution of network outputs over this interval (dashed grey ellipsoids) and the target output distribution that maximizes information transmission (solid black circle). **d)** Information-maximizing filters are learned with Independent Component Analysis [52]. Global filters that maximize information transmission for all image patches (gray frame) differ from context-optimal filters (colored frames) that maximize information transmission for patches originating from individual contexts. **e)** Biasing the code toward reducing the mismatched error rate yields a tradeoff between mismatched and matched error. Solid lines mark contours of constant global error. The optimal code that minimizes global error is marked with a star. **f)** Biased codes reduce the duration of mismatch, measured via the classification error per stimulus sample. Increasing the bias beyond the optimum results in a dramatic improvement of classification performance (main panel). This is accompanied by an abrupt change in the shape of receptive field (upper row; shown for context 1. Red barbells mark the maximum and minimum values of the receptive fields and are shown purely for visualization). **g)** Biased codes achieve performance intermediate between the static and oracle codes. The optimal code (*α*_NL_ = 0.14) achieves three-fold lower error than the naively-adaptive code (*α*_NL_ = 0).

As before, we consider an environment that can switch between these four contexts with a small but fixed probability per time. At each time point, an image patch *s*_*t*_ is sampled from the context specified by *c*_*t*_, encoded in the joint response 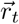 of the two neurons, and used to update a maximum likelihood estimate *ĉ*_*t*_ of the current context. This estimate is then used to adapt the receptive fields, specified by the encoding parameters *ψ*, for the task of maximizing information transmission over a window of time Δ*T*. In the low-noise limit, information transmission will be high when each neuron saturates the variance of its output, and neural activity is decorrelated [52, 57]. We can thus measure the task error at time *t* using the symmetrized Kullback-Leibler (KL) divergence between this ideal output distribution and the actual output distribution, computed using neural activity over the past Δ*T* time points (Fig 7c; Methods). We use the symmetrized, rather than non-symmetrized, KL-divergence because it conceptually matches the other (symmetric) error measures discussed here. We then average this error over time and context switches to compute the global KL error.

The set of context-optimal receptive fields 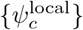 that maximize information about patches within each context *c* differ from one another (Fig 7d, lower rows), and from the set of global receptive fields *ψ*^global^ that maximize information about patches uniformly sampled from all contexts (Fig 7d, upper row). Such adaptation of receptive fields to high-order stimulus statistics has been observed in primary visual cortex but typically evokes more subtle effects than those generated by this simplified model [12, 21, 58]. We construct a naively-adaptive code that uses its context estimate to select among the context-optimal receptive fields. We then bias this code toward nonlocal task performance by minimizing the mismatched error rate. We do this by learning a set of receptive fields 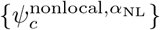 that maximize information about a set of image patches in which a fraction (1 − *α*_NL_) is sampled from the current context, and a fraction *α*_NL_ is sampled from the remaining contexts. We empirically found this biasing strategy to yield the largest reduction in global error, and so we focus on this strategy for the remainder of the section.

As in our previous examples, we find that stronger biases lead to lower mismatched error but higher matched error (Fig 7e). As a result, the optimal performance that minimizes global error is achieved for an intermediate bias of *α*_NL_ = 0.14 (starred points in Fig 7e-g). Interestingly, however, the duration of mismatch decreases dramatically for a small increase in bias beyond the optimal value (Fig 7f). While this small increase leads to slightly higher global error than the optimal code (Fig 7g), it is still well below the error attained by the naively-adaptive code. A neural population that needs to balance adaptation speed and information transmission could thus benefit from this tradeoff by greatly reducing the duration of mismatches at the cost of a small increase in global error. These significant improvements in performance are achieved by relatively subtle modifications of receptive fields (Fig 7f, top row). Taken together, this simple model illustrates the benefits of biasing the joint encoding of high-dimensional stimuli to improve global performance.

## DISCUSSION

Adaptive phenomena in neural coding have been a subject of experimental and theoretical research since Adrian’s and Zotterman’s discovery of adaptation [1]. The dynamics of adaptation have traditionally been characterized in terms of changes in experimentally-observable quantities—such as firing rates, tuning curves, and receptive fields—in response to changes in the statistics of incoming sensory stimuli [5, 9, 14, 24, 26, 59–64]. A large body of modeling work has focused on reproducing these dynamic changes [49, 65], or on simulating their mechanistic origins [16, 47, 66]. Other studies, both theoretical [67, 68] and experimental [7, 69], have examined the impact of adaptation dynamics on neural coding and perception. Together, these approaches can capture the dynamics of adaptation and explain their impact on specific computations. However, they do not specify the computations that would warrant such dynamics, nor can they explain why a particular set of dynamics might be more or less beneficial to an organism. Such questions fall within the domain of normative approaches to neural coding.

Existing normative accounts of sensory adaptation have largely been developed within the framework of the efficient coding hypothesis [27, 70]. Efficient coding studies traditionally derive optimal solutions for maximizing utility (e.g. [15, 35, 52, 57]) or minimizing error (e.g. [30, 43, 71–73]) in representing natural sensory stimuli under the assumption that their distribution is stable [15, 42, 74]. Natural environments, however, are constantly changing [55, 56, 75, 76], which is reflected in the dynamic and adaptive structure of sensory codes in the brain [10, 33, 77]. To account for this, efficient coding posits that adaptation serves to readjust the sensory code in order to maintain a maximal rate of information transmission [10–12, 26, 78], supporting the idea that adaptation reflects a dynamic reallocation of limited metabolic resources that serves to maximize performance within different environmental contexts. While these studies provide a normative objective for the outcome of adaptation, they do not explain the dynamic structure of resource reallocation that would best achieve this outcome, nor do they explain how this reallocation should optimally unfold in nonstationary environments. As a consequence, they cannot account for the diversity of adaptation dynamics that have been observed experimentally.

Here, we addressed this gap by extending classical theories of efficient coding to the study of adaptive neural computation in non-stationary environments. To our knowledge, this is the first extension of the efficient coding framework to nonstationary settings (however, see [79]), and the first normative framework that can account for the dynamics of adaptation. Our key insight is that in order to reliably and efficiently encode task-relevant information, sensory systems often need to rely on different solutions when the statistics of the environment are changing than when they are stationary. Such optimal performance requires that sensory systems not only accurately encode incoming stimuli when their distribution is constant in time, but also detect and adapt to changes in this distribution [11]. To rapidly infer such changes, neurons must accurately encode stimulus features that distinguish between different stimulus distributions. At any point in time, the most discriminative features are determined by the observer’s dynamically evolving belief about the stimulus distribution. The online estimation of changes in stimulus distributions from limited and noisy stimulus samples is thus an inherently complex problem, and is further complicated if the representation of these stimuli is lossy [80]. As a result, there will necessarily be periods of time when the system incorrectly estimates the stimulus distribution and the code is maladapted. Such periods of mismatch will result in a suboptimal use of resources—as has been observed experimentally via a drop in information rate immediately following a change in stimulus statistics [10, 11, 81, 82]—and consequently larger coding errors than would be achieved in periods of stationarity. Here, we provide a set of rules for deriving sensory codes that balance performance with the competing demands of estimating changes in the stimulus distribution and combating the adversarial effects of mismatches between this estimate and the stimulus distribution.

To achieve this balance, we considered a sensory system that uses the output of an adaptive encoder to construct an internal estimate of the current sensory context, and in turn uses this estimate to dynamically adapt the encoder. It has previously been suggested that adaptation is guided by an internal model that integrates new information over time [23, 31, 45], and several lines of work have shown that the relative timescales of adaptation are consistent with the predictions of ideal observer models [9, 24, 31, 47]. Here, we formalized the relationship between the encoder and the internal model of context, and we identified coding strategies that could optimally exploit this context knowledge in order to improve performance on a sensory task. We considered two different performance measures: minimizing decoding error in individual neurons, and maximizing information transmission in a small neural network. Regardless of the performance measure, we found that even subtle differences in sensory representations when the environment is stable can lead to dramatically different dynamics and task performance when the environment changes. Importantly, in all cases considered here, the optimal code was not the one designed to maximize task performance given a local context estimate, but rather was a code designed to simultaneously balance errors that arise when this estimate is inaccurate.

We then showed how this general theoretical setting can be used to derive quantitative predictions about neural dynamics—such as dynamic changes in firing rates and nonlinearities in the retina—that are comparable to phenomenological and mechanistic models of adaptation [16, 49, 65]. However, because these predictions are derived directly from a normative theory, our framework can additionally explain the putative computational purpose of these dynamics. To this end, we provided the first normative account of sensitizing and adapting dynamics in retina, which arise in our framework from two different optimality criteria: minimizing the average error in encoding stimuli, and maximizing the speed of adapting to changes in the distribution of these stimuli. Because this adaptation is guided by an internal estimate of the changing sensory context, its timescale varies nearly linearly with the volatility of the environment, a finding that has been observed in many different systems [10, 45, 49]. Our approach can therefore simultaneously account for qualitative features of firing rate dynamics during adaptation, and for statistical properties of such dynamics derived from ideal-observer models. To our knowledge, this is the first approach capable of synthesizing these different perspectives on adaptation.

The codes that produced these adapting- and sensitizing-like dynamics correspond to two solutions within a family of different codes. At one extreme, we identified a code that favors fine discrimination of stimuli within the current context at the cost of resolving stimuli that would signal a change in context (Fig 6a-b, left). At the other extreme, we identified a code that favors detecting changes over making fine discriminations (Fig 6a-b, right). In both cases, adaptation serves to dynamically reallocate a finite partitioning of the stimulus space in order to support these objectives in the new context (i.e., after adaptation). Previous work has shown that adaptation can indeed mediate tradeoffs between detection and discrimination at both a neural and behavioral level [7]. Here, we showed how such a shift should evolve over time, how it might be signaled by specific patterns of neural dynamics, and how these two different objectives might optimally be balanced (Fig 6a-b, center). Because we used decoding error to evaluate performance, the optimal solution that we derived ultimately favors local discrimination, and uses detection only as a means of improving discrimination. Other systems might instead favor the detection of changes in context over the discrimination of local stimulus details [80].

The distinctly different firing rate profiles of adapting and sensitizing cells have been observed across several different organisms [16, 48]. The fact that many systems exhibit these two distinct types of dynamics suggests that it might be beneficial for a population to perform change detection and fine discrimination within two distinct cell types. However, different tasks could warrant additional performance criteria; for example, in tasks such as predator localization or visual speed estimation, it might be more critical to stay below a maximum error threshold than to minimize global error. While these different objectives are handled separately within our current framework, it might be possible to derive coding schemes that simultaneously optimize multiple objectives. The question of how such criteria should be balanced within a population [83–85], and how these different objectives might relate to different functional cell types [86, 87], or to different stimulus features encoded by a single neuron [51], is the subject of future work.

Within the framework developed here, performance is determined by the interplay between a system’s ability to accurately encode incoming stimuli, and its ability to accurately detect changes in the underlying distribution of these stimuli. The precise structure of this interplay is scenario-specific and depends on multiple factors, including the statistics of the environment, the sensory task, and the nature of resource limitations within the sensory system. In some scenarios, achieving the optimal balance of errors between periods of match and mismatch might be computationally complex and difficult to instantiate in a biological system. However, we demonstrated that even simple, approximative coding strategies—such as linear interpolation of coding parameters—can improve global performance by biasing away from the naively-adaptive solution.

In all of the cases that we examined, such improvements in global performance necessitated a compromise between adaptation speed and task performance. These compromises are reminiscent of physical tradeoffs between the speed, energy, and accuracy of adaptation observed in biological circuits with negative feedback [88]. Understanding such tradeoffs from a normative perspective would require optimizing not only task performance and speed, but also instantaneous energy use. This could be viewed in terms of balancing additional constraints and objectives—in this case, minimizing energy and maximizing accuracy—with the need to keep up with a changing environment. The framework developed here could be generalized to such scenarios and used to identify the optimal interplay between multiple objectives of task performance and energetic cost.

Our approach relies on multiple idealizations; these include the use of encoders that can adapt within a single timestep, ideal observer models that optimally estimate changes in context, and an entropy-coding scheme that optimally transforms the output of the encoder to reduce firing rates. In practice, these idealized computations could be approximated by simpler computational schemes [77, 89, 90]. We also assumed that our model sensory systems could distinguish between a fixed number of discrete contexts. In practice, the number and statistical structure of relevant contexts depends on the statistical variability of natural scenes, which has itself been the subject of extensive research (e.g. [55, 56, 91–94]). Moreover, the number of distinct contexts to which a system adapts can itself be considered a resource constraint. Thus, under some scenarios, the cost of maintaining an adaptive code might exceed its benefits [95]. In such cases, it might be more beneficial to devote separate circuits to handling different contexts, or to use a single static code adapted to a set of contexts. Together, these scenarios can be formalized and compared within the current framework, and as such serve an avenue for future work.

### Outlook

To date, theories of efficient sensory coding have largely been concerned with stationary signals. By extending these theories to nonstationary environments, this work enables a normative description of a broad range of phenomena in neural coding that were previously outside of the domain of the efficient coding framework. A rigorous connection between the framework developed here and experimental data will require the development of statistical models that can capture time-varying changes in neural nonlinearities and receptive fields [22, 96, 97], and mathematical frameworks that can be used to study optimal coding in nonstationary environments from an information-theoretic perspective [98, 99]. Together, these developments may shed light on how adaptability and efficiency together guide the evolutionary design of neural representations, and thereby provide a normative understanding of a broad range of dynamic phenomena in adaptive neural coding.

## ACKNOWLEDGEMENTS

We thank David Kastner and Thomas Münch for generously providing figures from their work. We also thank Vivek Jayaraman, Marcella Noorman, and Tzuhsuan Ma for useful discussions and feedback on the manuscript. WM was supported by IST Plus and Marie Skłodowska-Curie Actions fellowship. AMH was supported by the Howard Hughes Medical Institute.

## SUPPLEMENTAL FIGURES

**Figure S1:**
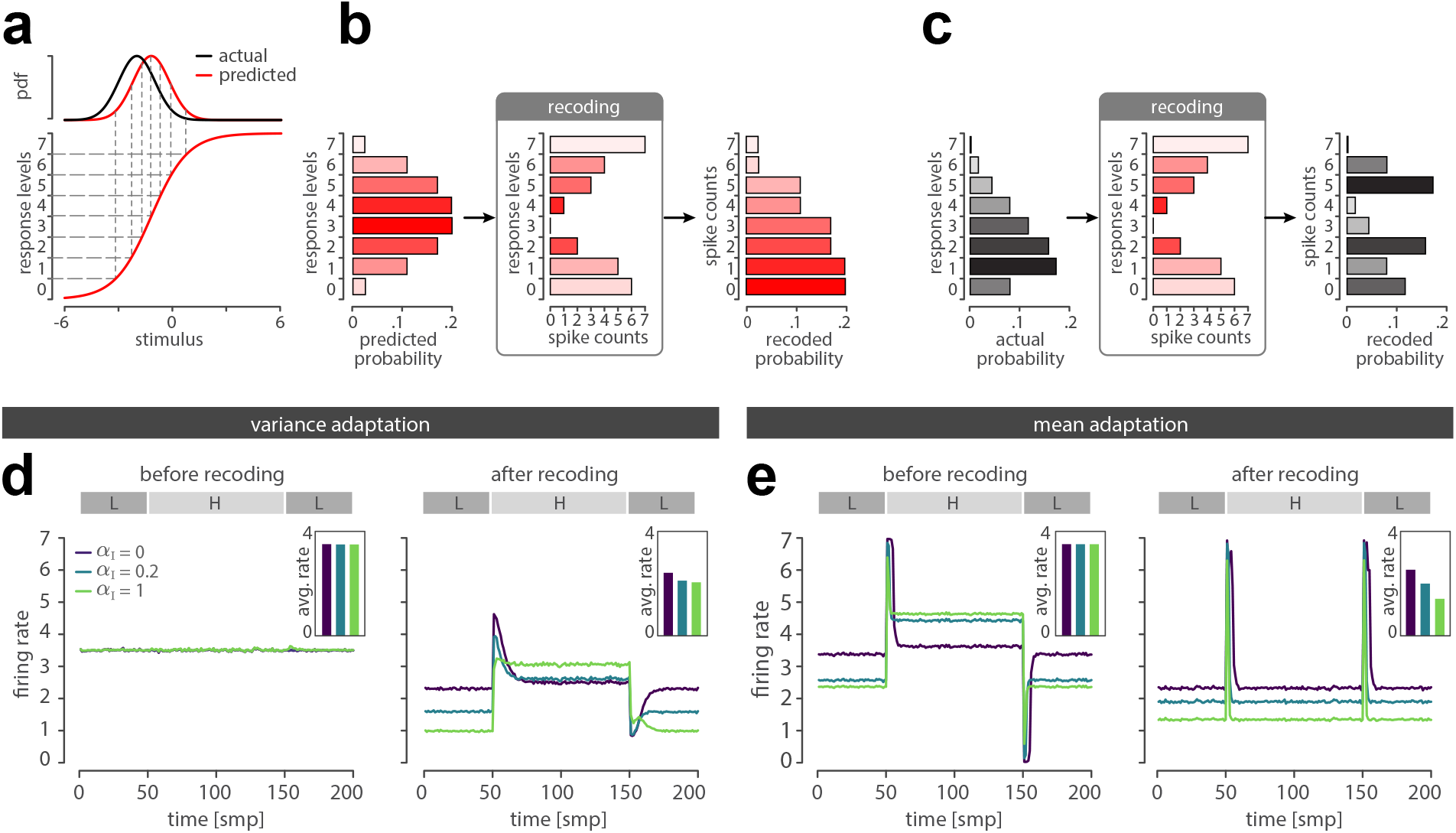
Entropy coding transforms response levels and reduces average firing rates. **a-c)** Entropy coding reassigns response levels to spike counts based on the predicted probability that a response level will be used. **a)** The encoding nonlinearity partitions stimuli drawn from the actual stimulus distribution (black) or predicted stimulus distribution (red) into discrete response levels. This partitioning determines the predicted (**b**, left) and actual (**c**, left) probability with which response levels will be used. **b)** Entropy coding reassigns spike counts in order of decreasing predicted probability (left); response levels that have higher predicted probability (darker red) will be assigned fewer spikes (middle). This leads to a recoded histogram that is weighted toward lower spike counts (right). The reassignment could be approximated by a quadratic nonlinearity, or by a thresholding exponential nonlinearity. **c)** The recoding scheme, which is based on *predicted* probability (**b**), determines how stimuli sampled from the *actual* stimulus distribution will be transformed into spike counts. **d)** Before recoding (left), firings rates do not change in response to a change between low (L) and high (H) stimulus variance, regardless of the value of *α*_I_. After recoding (right), all codes show a transient response to both increases and decreases in variance, and average firing rates over time are lower (inset). **e)** Before recoding (left), firing rates increase following a change from low (L) to high (H) stimulus mean, and decrease following a change from high to low mean. After recoding (right), all codes show a symmetric response to increases and decreases in mean, and time-averaged firing rates are lower (inset).

**Figure S2:**
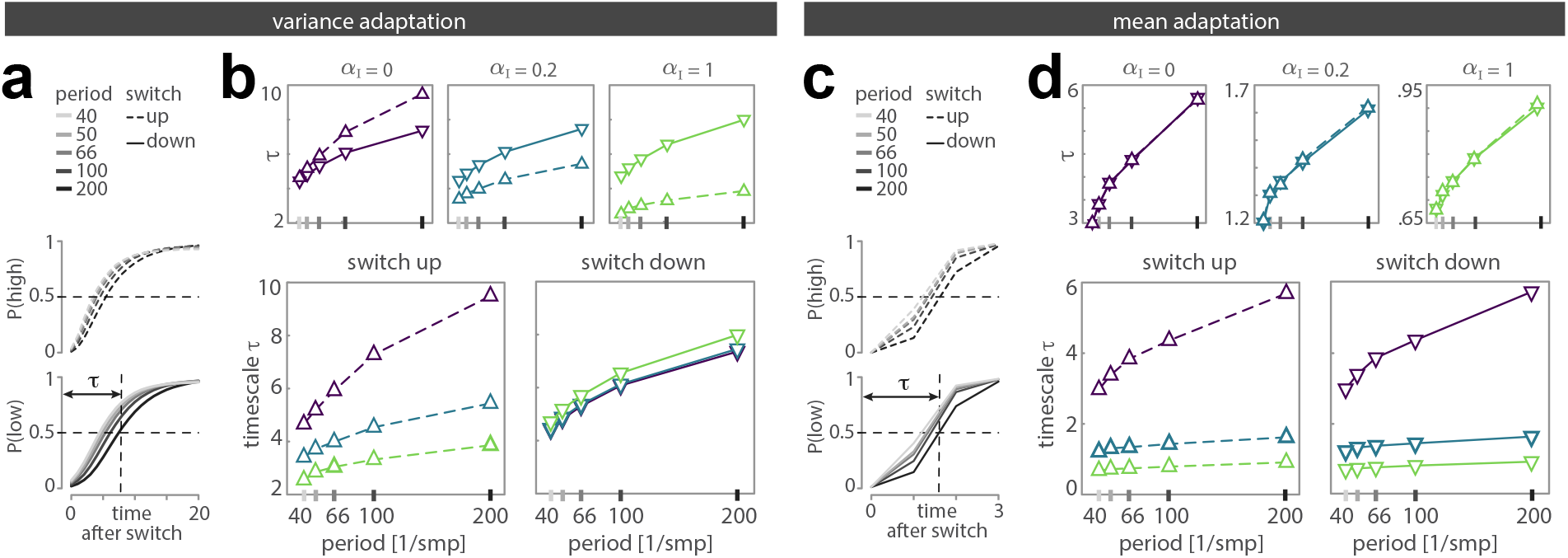
Volatility shapes posterior dynamics. **a**,**c)** As volatility increases (i.e., as the switching period decreases), the posterior is faster to respond to both increases (upper) and decreases (lower) in stimulus variance (**a**) and mean (**c**). We estimated this timescale by linearly interpolating the posterior before computing the time at which it crosses 0.5. **b)** When stimulus variance is changing in time, the naively-adaptive code (*α*_I_ = 0, upper left) is faster to respond to decreases in variance (solid lines) than to increases (dashed lines). In contrast, biased codes (upper middle, right) are faster to respond to increases in variance. This is largely driven by a more rapid response to increases in variance with increasing bias (lower left), but is also weakly affected by a slower response to decreases in variance (lower right). **d)** The posterior is equally fast to respond to increases and decreases in the stimulus mean (upper row), with a timescale that decreases as bias increases (lower row).

**Figure S3:**
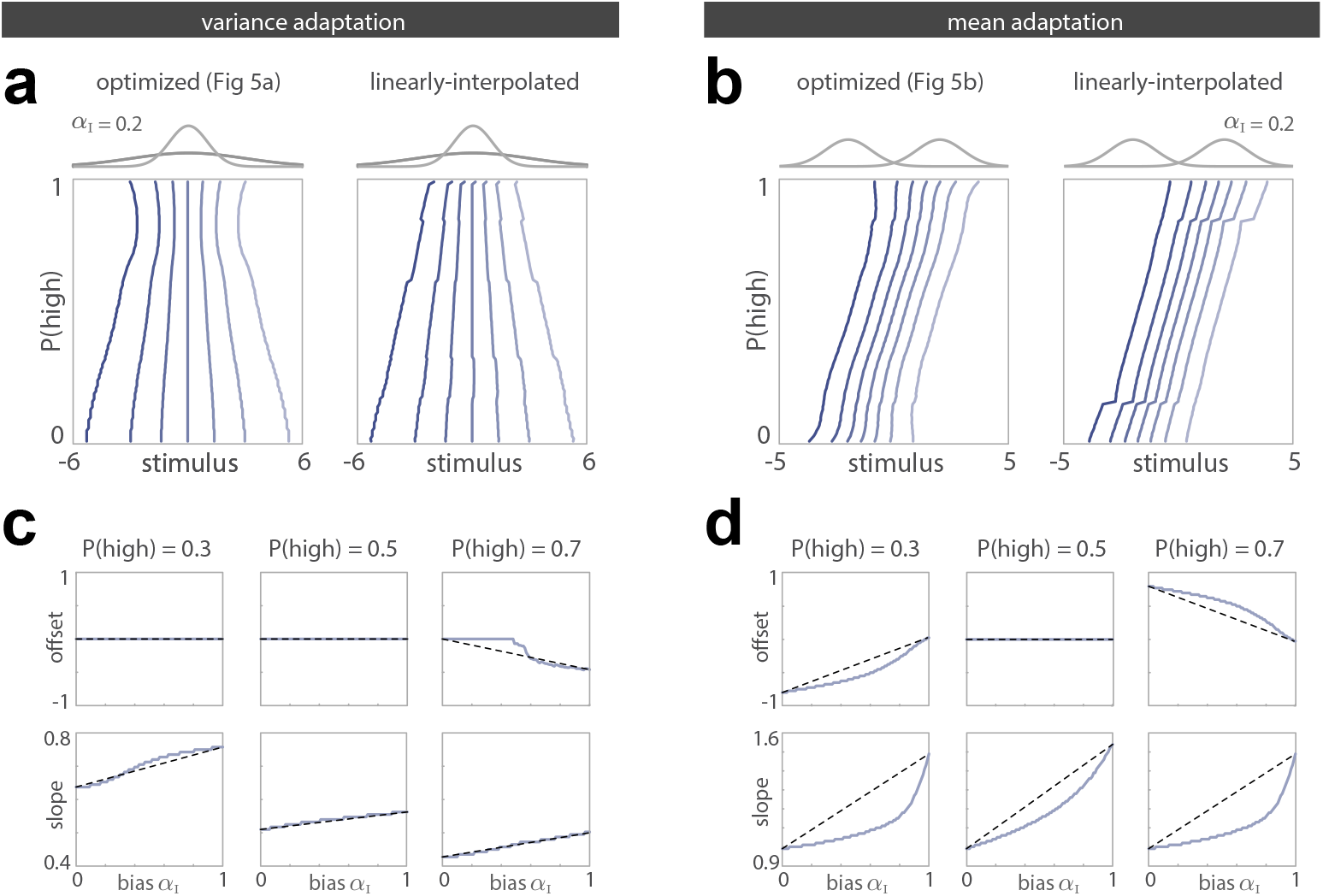
Optimal dynamics are robust to parameter variations. **a, b)** Optimal nonlinearies (left; shown in Fig 6a,b) closely resemble those derived by linearly-interpolating parameters between the naively-adaptive code and the fully-biased code (right). Shown here as a function of the system’s posterior belief 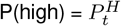, for variance adaptation (**a**) and mean adaptation (**b**). **c, d)** The dependence of the offset (upper row) and slope (lower row) of the optimal nonlinearities on the bias *α*_I_ can be approximated by a line that interpolates between the naively-adaptive code (*α*_I_ = 0) and the fully-based code (*α*_I_ = 1). Shown here for variance adaptation (**c**) and mean adaptation (**d)**, for several different values of the posterior belief P(high). **e, f)** Recoded firing rate dynamics vary smoothly as a function of *α*_I_ in response to changes in variance (**e**) and mean (**f**). Insets show the dynamics in response to switches from the low to high context (upper inset) or the high to low context (lower inset).

## METHODS

### Local contrast and luminance adaptation in a linear-nonlinear neuron

#### Adaptive encoder

We modeled a classical linear-nonlinear Poisson neuron that encodes incoming image patches in neural responses. This model has been broadly used to model stimulus-response properties in a range of sensory systems [36]. Here, this model neuron encodes incoming images patches of size 12 *×* 12 pixels by projecting them onto an oriented linear filter *ϕ*. The filter was randomly selected from a set of 64 filters learned with logistic Independent Component Analysis [52] from a dataset of images consisting of natural movie frames provided by [54]. The specific filter that we used is shown in Fig 1a; the choice of this filter did not qualitatively affect the results described here. The output of the filter *s*_*t*_ was mapped onto firing rate *λ*_*t*_ via an adaptive logistic nonlinearity:

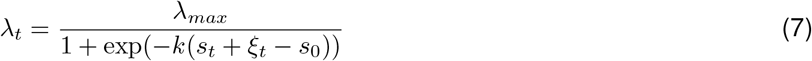

where *λ*_*max*_ is the maximal firing rate, *ψ* = {*k, s*_0_} are adaptive encoding parameters that specify the slope and offset of the nonlinearity, respectively, and 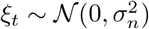 is an additive Gaussian noise with variance 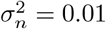. The discrete spike-count response of the neuron *r*_*t*_ was drawn from a Poisson distribution parameterized by rate *λ*_*t*_Δ*T*. For simplicity, we assume the time interval of a response to be equal to 1 (i.e., Δ*T* = 1).

#### Model environment

The encoder adapts by modifying the parameters of the encoding nonlinearity given a fixed filter *φ*. We therefore constructed contexts that capture the variability in its inputs (i.e., in the filter outputs) observed across an ensemble of natural scenes. We considered an environment that could switch between nine such contexts at a small but fixed probability per unit time, *p*_*s*_ (i.e., 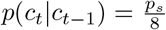 if *c*_*t*_ ≠ *c*_*t*−1_, and (1 − *p*_*s*_) otherwise). All results were generated with *p*_*s*_ = 0.01.

To mimic the variability in filter outputs that would be generated by visual exploration of a scene, we simulated saccadic trajectories across different regions of natural images. For a given trajectory, the distribution of filter outputs was well-approximated by a Gaussian; we thus used the set of trajectories to define the means and variances of a set of Gaussian contexts that spanned the range of filter outputs generated during this exploration. To generate the trajectories, we used the same dataset of images described above, and standardized each image in this dataset to have zero mean and unit variance. Each trajectory was generated via a Gaussian random walk and was used to extract a sequences of 10^3^ image patches of size 12 *×* 12 pixels; i.e., at each time step, the center of the image patch was moved by *n*_*x*_ and *n*_*y*_ pixels in the horizontal and vertical directions, respectively, where *n*_*x*_, *n*_*y*_ were drawn from a Gaussian distribution with zero mean and unit variance and rounded to the nearest integer. We randomly initialized the starting location of each trajectory, and we simulated 5 *×* 10^4^ trajectories in total. Three such trajectories are shown in the upper panel of Fig 1b. We then filtered the images patches within each trajectory using the oriented filter *ϕ*, which produced a single distribution of filter outputs for each trajectory (illustrated in the lower panels of Fig 1b for the corresponding trajectories). We then empirically estimated the mean and variance of each trajectory-specific distribution of filter outputs, thereby generating a distribution of means and variances across trajectories. We then selected 3 variance values 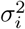 (0:6; 2:9; 13:6) that logarithmically spanned the range between the 10th and 90th percentile of the distribution of variances, and 3 mean values *µ*_*j*_ (− 2.3, 0, 2.4) that linearly spanned the range between the 1st and 99th percentile of the distribution of means. We then used all 3 × 3 = 9 combinations of means and variances to define a set of contexts *c* = {1, …, 9}, where each of the context labels specifies a pair of variance and mean indices *i*_*c*_, *j*_*c*_. Each sensory context thus defines a Gaussian distribution of filter outputs with variance 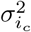 and mean 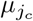, such that 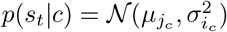. These distributions are shown in Fig 1c.

#### Sensory task

We evaluated the performance of the system by measuring the average error in decoding stimuli, ⟨(*s*−*ŝ*)^2^⟩. To compute this error, we decoded an estimate of the filter outputs using maximum-likelihood (ML) decoding [37]. At a given time *t*, we took the ML estimate *ŝ*_*t*_ to be the value that maximizes the probability of observing the response *r*_*t*_ given the encoding parameters *k, s*_0_. To numerically simplify the decoding process, we discretized the range of decoded filter outputs into 2^14^ values uniformly spaced between −16 and 16.

#### Observer

To aid in task performance, the system constructs an internal estimate of context *ĉ*_*t*_ that is used to select the appropriate encoding parameters at each timestep. We used a maximum-likelihood estimation scheme to infer the context identity *ĉ* from the neural response *r*_*t*_. For a given nonlinearity parametrized by its slope *k* and offset *s*_0_, and for a given context *c* parametrized by mean 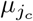 and variance 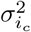, we empirically estimated the distribution of spike counts *p*(*r*_*t*_|*k, s*_0_, *c*). Over a window of time Δ*T*, the log-likelihood of responses *r*_*t*_ is given by:

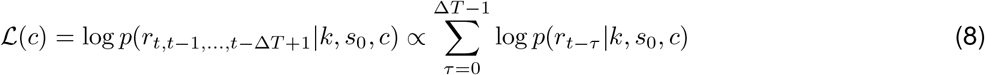

The ML estimate of the current context *ĉ* was found by iterating over all context labels *c*, and returning those that maximize the log-likelihood of the past Δ*T* responses:

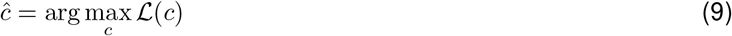

We set the history length Δ*T* to 20 steps, which corresponds to one fifth of the average duration of each context given the switching probability *p*_*s*_ = 0.01. For notational simplicity, we refer to *ĉ* as the context estimate, and will write *ĉ* ≠ *c* if the context estimate was correct (i.e., equal to the true context *c*), and *ĉ* = *c* if the context estimate was incorrect.

#### Optimization of nonlinear response functions

The set of encoding parameters Ψ = {*ψ*_*c*_} was optimized offline for each context *c*. During online simulations (described below), a single set of encoding parameters was selected from this set at each time-step *t* given the system’s estimate of context at time *ĉ*_*t*_.

We first determined the encoding parameters 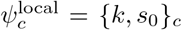 that minimize decoding error in each context *c*; these parameters specify the “context-optimal” nonlinearities shown in Fig 1c. We performed an exhaustive grid search over a 32 *×* 32 grid of parameter values, spanning the ranges [0.5, 1] for the slope *k*, and [−4, 4] for the offset *s*_0_. We generated 2 *×* 10^4^ stimulus samples *s* from the Gaussian distribution of filter outputs for each context. We then passed this set of stimulus samples through the encoding nonlinearity, defined by a given combination of slope *k* and offset *s*_0_, to generate the corresponding firing rates *λ*. We then randomly sampled a spike count *r* from a Poisson distribution parameterized by the rates *λ*. We used the ML decoding procedure described above to decode each stimulus and thereby obtain stimulus estimates *ŝ*. For each parameter combination, we computed the decoding error ⟨(*s ŝ*)^2^⟩ _*p*(*s*|*c*)_ averaged over a given context-specific distribution of stimuli. For each context *c*, we selected the combination of slope *k*^*c*^ and offset 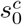 that minimized this average error.

An analogous procedure was used to determine the parameters *ψ*^global^ of the static nonlinearity shown in Fig 1c. Here, we uniformly sampled 2 *×* 10^4^ stimulus values from all nine contexts. The same grid search was used to determine the combination of slope and offset that minimize the decoding error ⟨ (*s* − *ŝ*)^2^⟩_*p*(*s*)_ averaged over the distribution of stimuli marginalized over contexts (i.e., 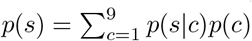, where 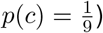).

In order to reduce either the mismatched error rate or the duration of mismatch, we biased each context-optimal nonlinearity towards a target nonlinearity. We did this by constructing convex combinations of context-optimal parameters 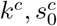, and fully-biased target parameters 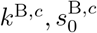, where B ∈ {I, NL} denotes whether the target parameters are optimized for inference or nonlocal task performance. We weighted these combinations by the coefficient *α*_B_ ∈ [0, 1]. For *m* ∈ {*k, s*_0_ }, we took 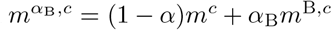. We used 50 values of *α* spaced evenly between 0 and 1 to construct two families of codes that were biased toward minimizing either the error rate or the duration of mismatches. As noted in the Results, this biasing strategy approximates the set of nonlinearity parameters by linearly interpolating between *k*^*c*^, 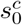 and target parameters *k*^B,*c*^, 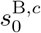. This approximative strategy, although not strictly optimal, results in a global error minimum that out-performs the naively-adaptive code. Moreover, in a simplified model of mean and variance adaptation (Fig 6), we found that a more principled, but more computationally exhaustive, biasing strategy can be well approximated by linear interpolation (see SI Fig S3a-d).

To bias codes toward reducing the duration of mismatches, we optimized a single target nonlinearity for context inference (shown in Fig 4c, right panel, *α*_I_ = 1). We again performed an exhaustive grid search over a 32 *×* 32 grid of parameter values, here spanning the ranges [0.1, 3] for the slope *k* and [− 4, 4] for the offset *s*_0_. We sampled 10^4^ stimulus values from each context, resulting in 9 *×* 10^4^ samples in total. We then generated corresponding firing rates, responses, and stimulus estimates as described above. For each individual neural response *r*, we then used the ML decoding procedure described above to construct an estimate of context *ĉ*; note that this is equivalent to maximizing the log-likelihood in Eq. 8 using a window of size Δ*T* = 1. We then selected the pair of nonlinearity parameters *k*^B^, 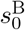 that maximized the probability *p*(*ĉ* = *c|k, s*_0_) of correctly classifying all nine contexts.

To bias codes toward reducing the mismatched error rate, we constructed a different target nonlinearity for each context (an example of which is shown in Fig 4c, middle, *α*_NL_ = 1). To minimize the mismatched error rate, we minimized the decoding error averaged across the distribution of stimuli encountered by the system at times when the system’s context estimate *ĉ* differs from the true context *c* (i.e., *c*′ ≠ *ĉ*). We denote this distribution as 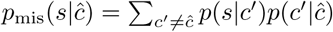.

The distribution *p*(*c*′ | *ĉ*) specifies the probability of mistaking the true context *c*′ for the estimated context *ĉ* ≠ *c*′. This distribution depends on the current estimate *ĉ*, since some contexts are easier to mistake for the current context than others. The pattern of these mistakes is summarized by the confusion matrix *M* (*ĉ, c*) = *p*(*ĉ, c*). The distribution *p*(*c* |*ĉ*), where *c* ≠ *ĉ*, can be obtained from the confusion matrix *M* by removing diagonal entries *p*(*ĉ, c* = *ĉ*) and renormalizing each row, such that 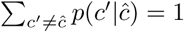.

We constructed a training dataset consistent with the distribution *p*_mis_(*s* | *ĉ*) for each context *ĉ* by running an online simulation (see below) with a naively-adaptive code for 3 *×* 10^5^ time steps. For each context, we then pooled together all stimulus samples that were incorrectly classified by the ML inference procedure (i.e., stimulus samples *s*_*t*_ for which the resulting context estimate *ĉ*_*t*_ was incorrect). We then performed an exhaustive grid search over a 32 *×* 32 grid of parameter values, here spanning the ranges [0.5, 1] for the slope *k* and [− 4, 4] for the offset *s*_0_. We then generated corresponding firing rates, responses, and stimulus estimates as described above, and we selected the pair of nonlinearity parameters that minimize the decoding error 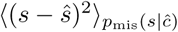.

#### Online simulations

We simulated dynamics over 3 *×* 10^5^ time steps. At each timestep *t*, the system maintained a context estimate *ĉ*_*t*−1_ from the previous timestep. This estimate was used to select the set of encoding parameters *ψ*_*t*_ = {*k, s*_0_ } to be used on the current timestep. We then randomly sampled a single stimulus sample *s*_*t*_ from *p*(*s*_*t*_ |*c*_*t*_). This stimulus sample was encoded in a response *r*_*t*_, which was then used to update the context estimate *ĉ*_*t*_. The response was also decoded to construct an estimate *ŝ*_*t*_, which was then used to compute the instantaneous decoding error (*s*_*t*_ − *ŝ*_*t*_)^2^.

To define periods of match and mismatch, we selected all time points for which *ĉ*_*t*_ = *c*_*t*_ and *ĉ*_*t*_ ≠ *c*_*t*_, respectively. The relative duration of mismatch, shown in Fig 4d and in the inset of Fig 4c, was computed by scaling the total number of mismatched time points by the total number of all time points. The matched error rate, shown in Fig 4d and in the inset of Fig 4c, was computed by averaging the instantaneous decoding error over all mismatched time points (i.e., 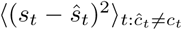). The product of these two terms was used to compute the mismatched error in Figs 4b and 4e. The matched error in Figs 4b and 4e was computed in an analogous manner, and the sum of matched and mismatched errors was used to compute the global errors in Figs 1d, 4b, and 4f. For visualization purposes, points in Fig 4 were smoothed using a boxcar filter of width 8, as indicated in the figure caption. This smoothing was done purely for display purposes; the optimal value of *α*_I_ and *α*_NL_ were selected prior to smoothing.

### Mean and variance adaptation in single neurons

#### Adaptive encoder

We modeled a single neuron whose response *r*_*t*_ to a stimulus *s*_*t*_ was determined by nonlinear function *f* (*s*_*t*_) = 1*/*(1 + exp(−*k*(*s*_*t*_ − *s*_0_))), where the adaptable encoding parameters *ψ* = {*k, s*_0_ } specify the slope and offset of the nonlinearity, respectively. We added Gaussian noise with standard deviation *σ*^2^ = 0.01 to the output of this nonlinearity. We then thresholded values below 0 or above 1, and we discretized the output into *n* response levels of equal width, ranging from 0 to *n* − 1. We took the discrete value at time *t* (representing a spike count) to be the neural response *r*_*t*_. All results were generated with *n* = 8 response levels.

#### Model environment

We considered an environment consisting of two contexts, a “low” context (*c* = *c*_*L*_) and a “high” context (*c* = *c*_*H*_). The environment could switch between these two contexts at a small but fixed probability per time, *p*_*s*_ (i.e., *p*(*c*_*t*_|*c*_*t*−1_) = *p*_*s*_ if *c*_*t*_ ≠ *c*_*t*−1_, and (1 − *p*_*s*_) otherwise). Unless otherwise specified, all results were generated with *p*_*s*_ = 0.01 (see below for details of simulations that vary this probability). We considered two environments, so-called ‘mean adaptation’ and ‘variance adaptation’, in which the context *c* parameterized either the mean or the standard deviation of a Gaussian stimulus distribution. In the former case, we took *p*(*s*|*c*) = 𝒩 (*s*; *c*, 1), with *c*_*L*_ = −2 and *c*_*H*_ = 2. In the latter case, we took *p*(*s*|*c*) = 𝒩 (*s*; 0, *c*^2^), with *c*_*L*_ = 1 and *c*_*H*_ = 3.

#### Sensory task

We evaluated the performance of the system by measuring the average error in decoding stimuli, ⟨ (*s* − *ŝ*)^2^⟩. To compute this error, we decoded an estimate of the stimulus *ŝ*_*t*_ = *p*_1_*r*_*t*_ + *p*_0_ using a linear function with an adaptable slope *p*_1_ and offset *p*_0_ (described in more detail below).

#### Observer

To aid in task performance, the system constructs an internal estimate of context *ĉ*_*t*_ that is used to select the appropriate encoding parameters, and also the appropriate decoding parameters, at each time step. We used an optimal Bayesian observer to construct the estimate *ĉ*_*t*_. To this end, the neural response *s*_*t*_ was first decoded as described above, and was then used to update the posterior distribution over *c*_*t*_. Because the context can only take two values, the posterior can be summarized by the probability that the environment is in the high context at time *t*, given by 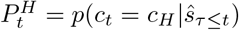. On each timestep, 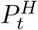 was updated with incoming stimulus estimates (as originally derived in [31] for raw stimulus values; also see [80]). We averaged the posterior to compute the point estimate 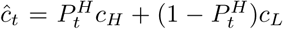, which could then be used to estimate the stimulus distribution *p*(*s*_*t*_|*ĉ*_*t*_) (where *p*(*s*|*ĉ*) = 𝒩 (*s*; *ĉ*, 1) in a mean-switching environment, and *p*(*s*|*ĉ*) = 𝒩 (*s*; 0, *ĉ*^2^) in a variance-switching environment). This estimate was fed back upstream and used to determine the optimal encoding and decoding parameters on the next timestep.

#### Optimization of nonlinear response functions

The set of encoding parameters Ψ was optimized offline for different values of the system’s current posterior 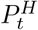. In practice, we discretized the posterior into 99 evenly spaced values between 0.01 and 0.99, and we determined the corresponding set of encoding parameters Ψ = {*ψ*_*n*_ }, where *n* ∈ [1, 99] indexes the possible values of the posterior (and corresponding values of *ĉ*). During online simulations (described below), a single set of encoding (and decoding) parameters was selected from this set given the system’s current estimate of context, *ĉ*.

For each value of the posterior (and each corresponding value of *ĉ*), we performed an exhaustive grid search over a 201 *×* 201 grid of parameter values, spanning the ranges [0, 2], [0, 1.5] for the slope *k* (in the mean- and variance-switching environments, respectively), and [−2.5, 2.5], [− 1, 0] for the offset *x*_0_ (mean- and variance-switching, respectively). We then generated 1 *×* 10^5^ stimulus samples drawn from *p*(*s*|*ĉ*). We passed this set of stimulus samples through the encoding nonlinearity, defined by a given combination of slope *k* and offset *s*_0_, to generate a distribution of spike counts *r*. We computed the parameters *p*_0_ and *p*_1_ of the optimal linear decoder that minimized the decoding error ⟨ (*s* − *ŝ*)^2^⟩_*p*(*s*|*ĉ*)_ between the true *s* and decoded *ŝ* stimulus values. The resulting set of decoded stimuli were used to update the estimate *ĉ*(*ŝ*) and compute the average inference error ⟨ (*ĉ*(*s*) − *ĉ*(*ŝ*))^2^⟩_*p*(*s*|*ĉ*)_. When iterated over all combinations of *s*_0_ and *k*, this procedure resulted in two error landscapes, 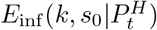 and 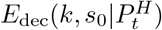, for each value of the posterior 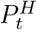.

In the mean-switching environment, the symmetry of the problem guarantees that the error landscapes are equivalent under the exchange *s*_0_ → (−*s*_0_) and 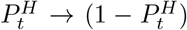. We used this equivalence to reduce numerical noise in our optimization through the following construction: 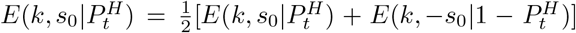. This was not performed for the variance-switching environment. We then normalized the values of each landscape to lie between 0 and 1, and we smoothed each landscape by replacing the error at a given entry (*s*_0_, *k*) with the average over a 12 *×* 12 region around the entry. We then constructed weighted combinations of the two landscapes: 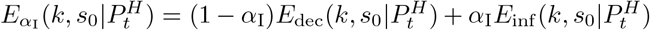, where *α*_I_ is a weighting factor that we varied between 0 (pure decoding) and 1 (pure inference). Finally, we found the combination of parameters (*s*_0_, *k*) (with corresponding decoding parameters (*p*_0_, *p*_1_)) that minimized the error 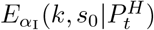 for each value of 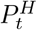 and each value of *α*_I_.

When *α*_I_ = 0, this procedure defines the parameters 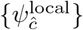 of the naively-adaptive code. When *α*_I_ *>* 0, this procedure defines the parameters 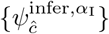 of a set of codes that are biased toward inference and thus toward reducing the duration of mismatch. We empirically found that this form of biasing led to the lowest global error. We used 101 values of *α*_I_ spaced evenly between 0 and 1. In the main text, results are shown for *α*_I_ = [0, .02, .04, .06, .08, .1, .2, .3, .4, .5, .6, .7, .8, .9, 1].

An analogous procedure was used to determine the parameters *ψ*^global^ and 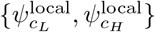 of the static and oracle codes, respectively. In the former case, we uniformly sampled 1 *×* 10^5^ stimulus values from both contexts (i.e., we sampled 5 *×* 10^4^ samples from each context). We used the grid search described above to determine the encoding and decoding parameters that minimize the decoding error ⟨ (*s* − *ŝ*)^2^⟩_*p*(*s*)_ averaged over the distribution of stimuli marginalized over both contexts (i.e., 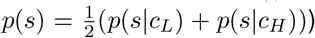). In the latter case, we sampled 1 *×* 10^5^ stimulus values from each context, and used the same grid search to determine the encoding and decoding parameters that separately minimize the decoding errors 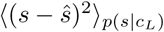 and 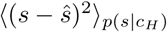.

#### Simulations

We simulated a probe environment that switched between *c*_*L*_ and *c*_*H*_ every 1*/p*_*s*_ = 100 timesteps; this probe signal is not unlikely given the generative process for *c*_*t*_ [31]. At each timestep *t*, the system maintained a posterior belief 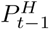 and corresponding point estimate *ĉ*_*t*−1_ from the previous timestep. The posterior belief specified the set of optimal encoding and decoding parameters to be used on the current timestep (as described above). We then randomly sampled a single stimulus sample *s*_*t*_ from *p*(*s*_*t*_|*c*_*t*_). This stimulus sample was encoded and decoded to construct an estimate *ŝ*_*t*_, which was then used to update the posterior belief and compute the instantaneous decoding error (*s*_*t*_ − *ŝ*_*t*_)^2^.

We defined a single trial of the probe environment to consist of 100 timesteps in the low state (*c*_*t*_ = *c*_*L*_), followed by 100 timesteps in the high state (*c*_*t*_ = *c*_*H*_). We simulated dynamics over 5000 continuous trials, and we averaged the results across trials. This resulted in trial-averaged values of the posterior, context estimate, and decoding error at each point in time. These trial-averaged values (shown in Fig 5b) were used to compute all of the remaining results in Fig 5.

To define periods of match, we selected all time points for which 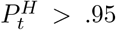 (when *c*_*t*_ = *c*_*H*_), and for which 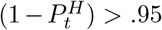 (when *c*_*t*_ = *c*_*L*_). All other timepoints were labeled as mismatched. The relative duration of mismatch, shown in Fig 5c, was computed by scaling the total number of mismatched timepoints by the total number of all timepoints (i.e., 200 timepoints). The mismatched error rate, also shown in Fig 5c, was computed by averaging the decoding error over all mismatched time points. The product of these two terms was then used to compute the mismatched error in Fig 5d. The matched error in Fig 5d was computed in an analogous manner, and the sum of matched and mismatched errors was used to compute the global error in Fig 5e.

We separately varied the volatility of context switches by varying the switching probability *p*_*s*_. All results in Figs 5b-e and 6a-d were generated using the probe signal described above, with *p*_*s*_ = 0.01. The results in Figs 5f-i and 6e-f were generated using a probe environment with *p*_*s*_ = [.005, .01, .015, .02, .025] (corresponding to a period of 1*/p*_*s*_ = [200, 100, 66, 50, 40]). For each value of *p*_*s*_, we simulated 5000 continuous trials of the corresponding probe environment, and averaged results across trials as described above. The impact of the switching probability on the duration, error rate, and error of matches and mismatches is shown in the upper panels of Fig 5g-i.

To compute the net change in duration, shown in the lower panels of Fig 5g, we subtracted the relative duration of the naively-adaptive code from all codes. We did this separately for each value of *p*_*s*_. To compute the percent change in the mismatched error rate, mismatched error, matched error, and global error (shown in the lower panels of Fig 5g-i), we first subtracted the corresponding error of the naively adaptive code from all codes, and we then scaled the resulting change in error by the difference in error between the static and oracle code.

The results shown in Fig 6a-b summarize the behavior of the optimal nonlinearities 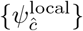 (for *α*_I_ = 0) and 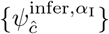 (for *α*_I_ *>* 0) for different values of 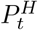 (and thus different values of *ĉ*). For each possible value of 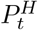 (as described above, we discretized the posterior into 99 values in total), we used the corresponding values of *k, s*_0_ to compute the encoding nonlinearity. We discretized the output of this nonlinearity into *n* = 8 discrete response levels (as described above), and we then determined the *n* − 1 values along the stimulus axis that, when passed through this nonlinearity, marked the transitions between these *n* response levels. Fig 6a-b shows these *n* − 1 stimulus values as a function of 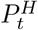.

#### Entropy coding

In Fig 6c-d, we used an entropy coding procedure to recode the discrete output of the encoding nonlinearity (also see SI Fig S1). Entropy coding assigns codewords to response levels based on the predicted probability that the response level will be used, such that shortest codewords (lowest spike counts) are assigned to the most probable levels [100]. In our case, we used entropy coding to reassign spike counts [0, …, *n* − 1] to the *n* response levels of the encoding nonlinearity based on the system’s current estimate of the stimulus distribution, *p*(*s*_*t*_ |*ĉ*_*t*−1_). We performed this recoding for each timepoint and on each individual trial. We first sampled 5 *×* 10^4^ stimulus samples from this estimate stimulus distribution. We then passed these stimulus samples through the encoding nonlinearity specified by this same context estimate, and we binned the output into the *n* response levels (SI Fig S1a). This resulted in a histogram of counts specified how often each response level was expected to be used (SI Fig S1b, left). The response level that had the highest count was assigned 0 spikes. The remaining response levels were sorted in descending order according to their count, and were assigned the remaining [1, …, *n* 1] spikes (SI Fig S1b, center). This resulted in a recoded histogram for which the fewest spikes are expected to be used with the greatest frequency (SI Fig S1b, right). We used then used this mapping to assign a spike count *r*_*t*_ to the stimulus sample *s*_*t*_ that was drawn at the given timepoint and on the given trial (SI Fig S1c). This resulted in a new recoded raster of spike counts that we used to compute the trial-averaged firing rates shown in the left columns of Fig 6c-d (SI Fig S1d-e shows the firing rates before and after recoding). We compared the firing rates in the left column of Fig 4c to data from adapting and sensitizing ganglion cells in the retina of salamander and mice, responding to step increases in contrast (figure reproduced from Fig 1a in [16]). We compared the firing rates in the left column of Fig 4d to data from two ON ganglion cells in mouse retina, both responding to the same two step increases in luminance (figure reproduced from Fig 4 in [46], using ND6 filters).

In Fig 6e-f, we computed the timescale of these firing rate dynamics following an increase in stimulus variance (Fig 6e) or mean (Fig 6f). We first performed the same entropy recoding described above to the simulations generated for different values of *p*_*s*_. We then fit an exponential of the form *a* exp(− *t/b*) + *c* to the resulting trial-averaged firing rates. To perform this fitting, we used the first 40 timepoints (equal to the period of the fastest context switches) following an increase in variance or mean. For large values of *α*_I_, there was a slower initial increase in firing rate after an increase in variance, followed by exponential decay. We thus first determined the time Δ*t* until the firing rate crossed a threshold of *BL*+0.5(*FR*_max_ − *BL*), where *FR*_max_ is the maximum firing rate, and *BL* is the baseline firing rate determined from the final 5 timepoints before a decrease in mean or variance. We removed timepoints *t <* Δ*t*, performed the exponential fitting on the remaining timepoints, and computed the total timescale *τ* = *b* + (Δ*t* − ⟨Δ*t* ⟩) (where ⟨Δ*t* ⟩ is averaged over different values of *p*_*s*_ for a given value of *α*_I_. The term (Δ*t* − ⟨Δ*t* ⟩) was only nonzero for large values of *α*_I_ in response to changes in variance; in all other cases, Δ*t* = 1 for all values of *p*_*s*_, and thus this term was 0. This procedure is qualitatively similar to the fitting used in [45] and shown in Fig 6e-f (reproduced from Figs 1e and 2e in [45]).

To summarize the behavior across different values of *α*_I_, we fit a line to the relationship between the timescale *τ* and the switching period, 1*/p*_*s*_. Examples of these best-fitting lines are shown in the insets of Fig 6e-f for different values of *α*_I_.

### Adaptation of receptive fields

#### Adaptive encoder

We modeled a population of two neurons that jointly encode image patches sampled from natural scenes. An incoming image patch *s*_*t*_ was transformed into a response vector 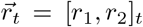 via a pair of adaptable receptive fields (implemented as adaptable linear filters). This response vector was then used both to perform the sensory task, and to update an internal estimate of context, as described below.

#### Model environment

We considered an environment consisting of four contexts *c*. The environment could switch between these four contexts at a small but fixed probability per time, *p*_*s*_ (i.e., 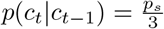 if *c*_*t*_ ≠ *c*_*t*−1_, and (1 − *p*_*s*_) otherwise). All results were generated with *p*_*s*_ = 0.01.

To define the set of contexts, we first sampled 2 *×* 10^5^ image patches of 12 *×* 12 pixels from standardized images taken from a natural movie dataset, as described above. We then performed Principal Component Analysis (PCA) on this set of images. We retained the first two first principal components, to which we applied a K-mean clustering algorithm in order to define four distinct clusters. We then used these clusters to assign individual image patches to one of these four contexts.

#### Sensory Task

We evaluated the performance of the system on the task of maximizing information transmission. As specified by the logistic ICA model [52], information maximization is achieved by obtaining maximally independent filter outputs with logistic mariginal distributions (see below).

To measure task error, we computed the distance between the observed distribution of filter outputs and the information-maximizing distribution of filter outputs over a time window Δ*T*. To simplify this computation, we summarized the observed distribution of filter outputs over the period [*t* − Δ*T* + 1, *t*] by a normal distribution *p*_*t*_(*r*_1_, *r*_2_) = 𝒩 (*µ*_*t*_, Σ_*t*_), where *µ*_*t*_ and Σ_*t*_ are the 2 *×* 1 vector of means, and 2 *×* 2 matrix of covariances, computed over the interval Δ*T*. Since the mean filter output does not affect information transmission, we subtracted this prior to evaluating the error. The Gaussian approximation of the information-maximizing distribution was defined as *p*^*G*^(*r*_1_, *r*_2_) = *N* (*µ*^*G*^; Σ^*G*^) where *µ*^*G*^ = [0, 0] is the vector of means, and Σ^*G*^ is the diagonal covariance matrix. The diagonal entries in Σ^*G*^ were equal to 3.29; i.e., the variance of the logistic distribution with a unitary scale parameter, as assumed by the ICA model [52] (see below). We quantified the distance between *p*_*t*_(*r*_1_, *r*_2_) and *p*^*G*^(*r*_1_, *r*_2_) using the symmetrized KL-divergence, *D*_*KL*_(*p*_*t*_ ‖ *p*^*G*^) + *D*_*KL*_(*p*^*G*^ ‖*p*_*t*_), and we used this KL error as our measure of task error.

#### Observer

To aid in task performance, the system constructs an internal estimate of the current context from the output of the receptive fields. To compute this estimate, we used a Gaussian Mixture Model (GMM) to approximate the distribution of filter outputs within each context. The parameters of the GMM were defined by the mean filter output within each context, and the covariance in filter outputs across all four contexts. The context estimate *ĉ* was taken to be the context that maximizes the likelihood of filter outputs given the parameters of the GMM.

#### Optimization of receptive fields

To obtain model receptive fields, we performed Independent Component Analysis (ICA) [52] on image patches belonging to each individual context. Since our goal was to learn a set of two receptive fields, we reduced the dimensionality of image patches to the first two principal components (as described above) prior to applying ICA. This resulted in a pair of context-optimal receptive fields for each of the four contexts. A single set of global receptive fields was obtained in the same manner, by performing ICA on the first two principal components of the ensemble of images patches across all four contexts.

We used ICA to learn context-specific filters that were biased towards minimizing the mismatched error rate. For each context, we created a training dataset in which (1 − *α*_NL_)*N* patches were sampled from the current context, and *α*_NL_*N* patches were sampled uniformly from all other contexts, where *N* = 10^4^ is the total number of patches. We then trained ICA filters on that new dataset. We considered 10 values of *α*_NL_ spaced linearly on the interval [0, 0.625]. We found empirically that this biasing strategy resulted in the largest decrease of the total KL-error with respect to the naively-adaptive code (*α*_NL_ = 0).

#### Simulations

We simulated dynamics over 2 *×* 10^5^ time steps. At each timestep *t*, the system maintained a context estimate *ĉ*_*t*−1_ from the previous timestep. This estimate was used to select the set of receptive fields to be used on the current timestep. We then randomly sampled a single image patch from the ensemble of patches specified by the current context *c*_*t*_. This image patch was encoded in a response *_r*_*t*_ as described above, which was then used to update the context estimate *ĉ*_*t*_. The KL error was computed from these responses every 10 timesteps over intervals of Δ*T* = 1*/p*_*s*_ = 100 timesteps. A given interval was defined to be “matched” if at least 80% the context estimates from that interval were correctly classified via the GMM inference process described above. Otherwise, the interval was defined to be “mismatched”. To compute the matched and mismatched errors shown in Fig 7e, we first summed the KL error per interval over all matched and mismatched intervals, respectively, and then scaled these by the total number of all intervals. We then summed these two errors to compute the global errors shown in Fig 7g. Classification error, shown in Fig 7f, was estimated by fitting the GMM to outputs of context-specific filters specified by each value of *α*_NL_ and computing the misclassification rate for individual stimulus samples (image patches).

### Scenario dependence of global error

#### Mathematical analysis

To better understand the dependence of global error on the bias coefficients *α*_I_ and *α*_NL_, we consider a simple, analytically-treatable setting.

Recall that the global error can be written as a function of the bias *α*_B_ ∈ [0, 1]:

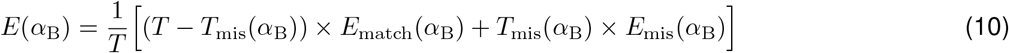

where B ∈ {I, NL} denotes biasing towards inference or nonlocal task performance, respectively. For simplicity, we now assume that each of these terms is a linear function of *α*_B_. These terms can thus be written as:

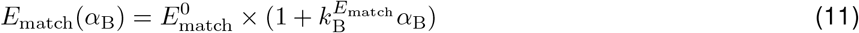

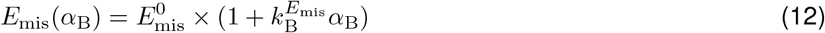

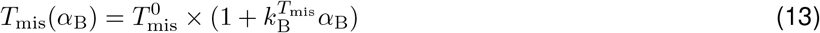

where 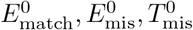 respectively denote the matched error rate, mismatched error rate, and duration of mismatch for naively adaptive code.

The linear coefficients 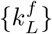 determine the rate at which each of these terms vary with *α*_*L*_. By construction, codes that vary with *α*_I_ and *α*_NL_ are designed to minimize *T*_mis_ and *E*_mis_, respectively. We can use this to constrain the sign of each coefficient, and the mutual relationship between coefficients. For codes biased toward inference, we can impose the following constraints on the dependence on *α*_I_:

- 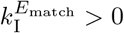: biased codes will necessarily increase the matched error rate
- 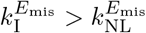: codes biased toward inference can either increase or decrease the mismatched error rate, but the rate of change must be larger for a code biased toward inference than one biased toward nonlocal task performance (since the latter was explicitly designed to minimize the mismatched error rate)
- 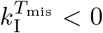 and 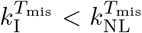: codes biased toward inference must decrease the duration of mismatch, and at a faster rate than the code biased toward nonlocal task performance (since the former was explicitly designed to minimize the duration of mismatches)

Analogously, for codes biased toward nonlocal task performance, we can impose the following constraints on the dependence on *α*_NL_:

- 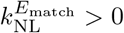: biased codes will necessarily increase the matched error rate
- 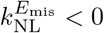 and 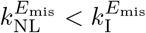: codes biased toward nonlocal task performance must decrease the mismatched error rate, and at a faster rate than the codes biased toward inference (since the former was explicitly designed to minimize mismatched error rate)
- 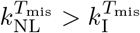: codes biased toward nonlocal task performance can either increase or decrease the duration of mismatches, but the rate of change must be larger for a code biased toward nonlocal task performance than one biased toward inference (since the latter was explicitly designed to minimize the duration of mismatches)

We can now substitute the *α*_B_-dependent error terms in Eq. 10 with their linear approximations in Eqs. 11-13. The global error then becomes a quadratic function of *α*_B_:

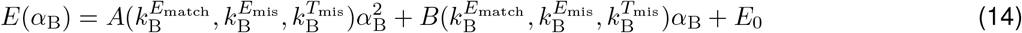

where *E*_0_ is the global error of the naively-adaptive code (*α* = 0). The coefficients of the quadratic equations have the following dependence on {*k*_B_}:

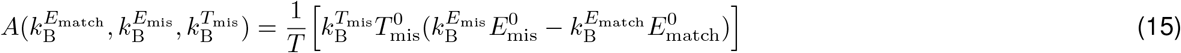

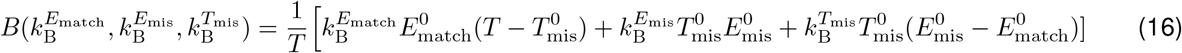

We use this quadratic approximation of global error to identify regimes in which one of the biasing strategies outperforms the other.

#### Visualizations

To illustrate the scenario dependence of global error, as shown in Fig 3, we used the following parameter values: 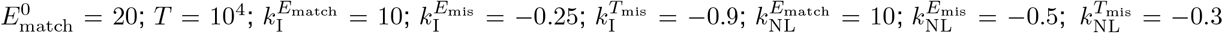. We first analyzed a single environment and determined the values of *α*_I_ and *α*_NL_ that produced the lowest global error (panel 3c); this environment was defined by 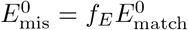 and *T*_mis_ = *f*_*T*_ *T*, where we chose *f*_*T*_ = 0.9 and *f*_*E*_ = 10. We then scanned across environments by varying *f*_*T*_ and *f*_*E*_; we sampled 100 evenly-spaced values of *f*_*T*_ between 0 and 0.99, and 1001 evenly-spaced values of *f*_*E*_ between 0 and 30. We first determined the values of *α*_I_ and *α*_NL_ that produced the lowest global error within the family of codes biased toward inference versus nonlocal task performance, respectively (panel 3d). We then determined which of the two families produced the lowest global error, and for what value of *α*_B_ (panel 3e). Finally, we varied 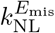 while keeping all other parameters fixed (using 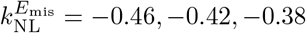), and separately varied 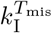 while keeping all other parameters fixed (using 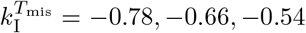). In all cases, the parameters satisfied the constraints listed above, and all errors were non-negative.

## Code availability

Custom code for parameter optimizations, simulations, and analysis was written in Matlab and is freely available upon request.

## Data availability

Data sharing not applicable to this article as no datasets were generated or analyzed during the current study.

